# DTH1-dependent remodeling of the lipid and protein environment of lipid droplets during nutrient recovery in Chlamydomonas

**DOI:** 10.64898/2026.10.06.757057

**Authors:** Carla Blot, Stéphan Cuiné, Bertrand Légeret, Marie Bertrand, Alwéna Tollec, Yohann Couté, Jean-Luc Putaux, Marina Siponen, Yonghua Li-Beisson

## Abstract

Lipid droplets (LDs) play a central role in cellular carbon and energy homeostasis, in particular through the roles played by numerous LD-associated proteins. A long-standing question concerns how these proteins are molecularly organized at the seemingly overcrowded LD surface. Delayed in TAG Hydrolysis 1 (DTH1) has been identified as a key protein for efficient LD degradation in *Chlamydomonas reinhardtii*, however the underlying molecular mechanism remains unknown. DTH1 contains several structural features – including a coiled-coil domain, a phosphatidylethanolamine (PE) specific lipid-binding domain, intrinsically disordered regions, and an α-helical C-terminal domain, that make it a compelling candidate for acting as a molecular scaffold in protein recruitment and/or organelle tethering. In this study, we used the Chlamydomonas knock-out mutant *dth1* as an entry point to begin dissecting the molecular organization of proteins at the LD surface. Comparative lipidomic and proteomic analyses of isolated LDs from both WT and *dth1* mutants revealed several key features associated with LD degradation during nutrient recovery. First, PE emerged as an important lipid species involved in LD degradation. Second, active communication occurs among multiple subcellular organelles particularly between LDs, peroxisomes and mitochondria. Third, DTH1 was found to play a critical role in remodeling both the LD proteome and lipidome. Furthermore, detailed analyses of LD degradation during nitrogen recovery uncovered the formation of micro-LDs, a phenomenon not yet reported in microalgae. Collectively, these results highlight the tight coordination of metabolic activities across subcellular compartments and emphasize the importance of intracellular carbon trafficking during stress recovery.

**Significance statement:** Lipid droplets are central hubs of cellular carbon storage and mobilization, yet how their protein and lipid composition is organized during turnover remains poorly understood. Using *Chlamydomonas reinhardtii*, we show that DTH1-dependent remodeling of the lipid droplet proteome and lipidome is associated with coordinated carbon mobilization involving peroxisomes and mitochondria, and identify micro-LD formation as a previously unreported feature of nutrient recovery.

## Introduction

Triacylglycerols (TAGs) are the universal form of carbon storage in living cells and are the primary constituents of oils used in food production, green chemistry and renewable fuels. In biological systems, when TAGs accumulate above a threshold concentration (>3-5 mol% locally), they are segregated from cellular membranes and packaged into lipid droplets (LDs) (Chorlay *et al*., 2019). In microalgae, LDs form in response to environmental stress and serve as dynamic carbon and energy reservoirs that can be mobilized through lipid catabolism when favorable conditions return (Li-Beisson *et al*., 2021). Reflecting their physiological importance and biotechnological potential, LDs have become a major focus of research across evolutionarily diverse microalgal lineages, including the model green microalga *Chlamydomonas reinhardtii* (hereafter Chlamydomonas)(Moellering and Benning, 2010; Goold *et al*., 2015; Tsai *et al*., 2015), the diatom *Phaeodactylum tricornutum* (Yoneda *et al*., 2016; Le Moigne *et al*., 2025; Murison *et al*., 2025), and stramenopiles such as *Nannochloropsis spp* (Vieler *et al*., 2012). Across all algal species examined to date, LDs are rapidly assembled and degraded in response to environmental cues. Moreover, mutants defective in LD turnover exhibit impaired recovery and delayed growth resumption following nitrogen (N) starvation (Kong *et al*., 2018; Lee *et al*., 2020). These observations emphasize the critical role of LD turnover in algal stress adaptation and point to the existence of tightly regulated mechanisms governing LD dynamics.

Unlike other subcellular organelles, LDs possess a distinctive “inside-out” architecture, consisting of a hydrophobic, neutral lipid core surrounded by a membrane lipid monolayer decorated with a diverse array of proteins (Moellering and Benning, 2010; Chapman *et al*., 2012; D’Andrea, 2016; Ischebeck *et al*., 2020). As a result, many LD functions are mediated by proteins present at their surface. Due to their low buoyant density, LDs can be efficiently isolated from other cellular components by sucrose density-gradient centrifugation, facilitating detailed proteomic and lipidomic analyses (Goold *et al*., 2015). Proteomics studies of purified LDs have revealed the presence of hundreds of associated proteins across diverse species, including structural proteins, enzymes involved in lipid metabolism, and numerous proteins of unknown functions or proteins not previously linked to LD biology (Moellering and Benning, 2010; Nguyen *et al*., 2011; Vieler *et al*., 2012; Goold *et al*., 2016; Yoneda *et al*., 2016). Despite the growing inventory, only a limited number of LD-associated proteins have been functionally characterized in Chlamydomonas (reviewed in Li-Beisson *et al*., 2021). These include the major lipid droplet protein (MLDP), the α/β-hydrolase domain containing protein 1 (ABHD1) (Torres-Romero *et al*., 2024), a long-chain acyl-CoA ligase (LCS2) (Bai *et al*., 2022), a diacylglycerol (DAG) lipase (LIP1) (Li *et al*., 2012) and a TAG lipase (LIP4) (Warakanont *et al*., 2019), despite the presence of the latter three proteins on LDs have not yet been experimentally validated by another means than LD proteomics.

In addition, forward genetic approaches have identified other factors involved in LD turnover, including the Delayed in TAG hydrolysis 1 (DTH1), a key regulator of LD turnover during recovery from N starvation (Lee *et al*., 2020). Furthermore, co-immunoprecipitation experiments using anti-MLDP antibodies provided evidence for the existence of supramolecular protein complexes at the LD surface in Chlamydomonas, suggesting that LD-associated proteins operate within highly organized molecular assemblies rather than isolated factors (Tsai *et al*., 2015). Given the central role of associated proteins in regulating LD dynamics, it is well established in many eukaryotic systems that LD proteomes are highly dynamic with proteins being recruited or released from LDs in response to changes in cellular metabolic and nutritional status (D’Andrea, 2016). In microalgae, however, our understanding of LD proteome remodeling remains limited, and a dynamic view of how LD-associated protein remodeling in response to environmental fluctuations is still lacking.

To identify the protein modules involved in LD turnover in Chlamydomonas, we used the LD-associated protein DTH1 as a molecular entry point to investigate the metabolic and cellular events underlying LD remobilization during recovery from N starvation. Although DTH1 has been established as a key regulator of LD turnover (Lee *et al*., 2020), the molecular mechanisms through which it promotes TAG mobilization remain largely unknown. Specifically, we sought to determine whether DTH1 contributes to the dismantling of the LD protein coat and/or lipid coat monolayer, whether it facilitates the recruitment of additional proteins to the LD surface to promote degradation, or whether it influences LD trafficking and interaction with other cellular organelles. To address these questions, we first characterized LD dynamics in wild-type cells during both TAG accumulation under N starvation and TAG degradation during recovery from nutrient depletion. Using an integrated approach combining lipidomics, proteomics and imaging analysis, we compared the composition, behavior, and remodeling of LDs from WT and *dth1* mutant strains to identify DTH1-dependent processes associated with LD turnover.

## Results

### Structural features of DTH1 protein by in silico analysis

To gain insight into the precise function of DTH1, in-silico analyses were conducted. Apart from its described role in LD degradation and the significance of the previously characterized “lipid-binding domain” (LBD), the remaining structural organization of DTH1 remained unknown. Taking advantage of the recent development in AlphaFold3 (Abramson *et al*., 2024) and other artificial intelligence-assisted tools, we performed an in-depth analysis of the DTH1 protein at the primary, secondary and tertiary levels. Although multiple alternative transcripts were predicted for this locus (Cre06.g261400), immunoblot analyses demonstrated that the mature DTH1 protein is likely of ∼157 kDa (Lee *et al*., 2020). We have therefore focused on this protein sequence for all subsequent analyses. A 3D structure of DTH1 predicted by AlphaFold3 is presented in **Figure 1A**. The most striking characteristic of DTH1 is its predominantly disordered nature, as predicted by PONDR (Xue *et al*., 2010) (**Figure 1B**).

**Figure 1.**
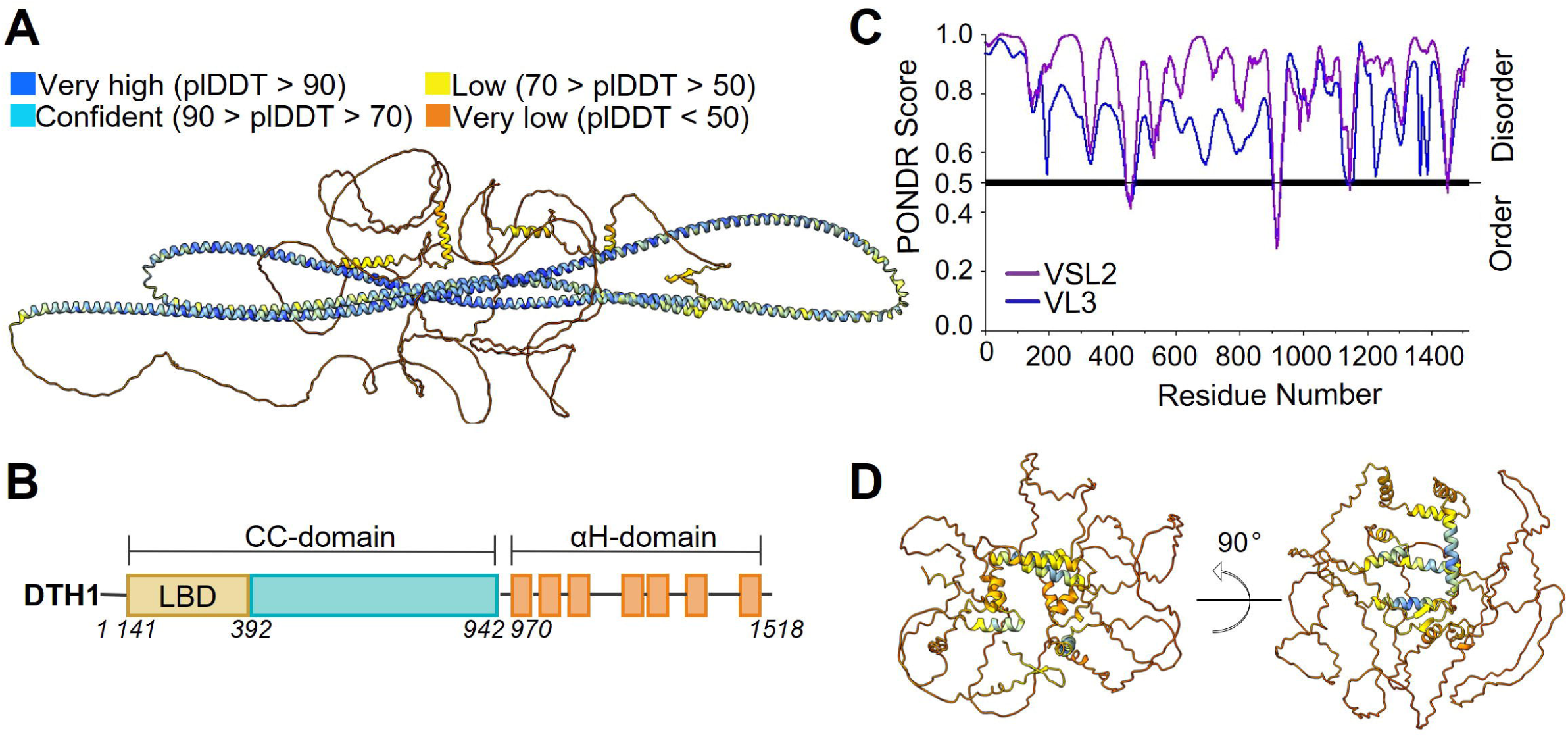
Predicted structural features of DTH1. **A.** AlphaFold3 structural prediction of the 157 kDa DTH1 protein, colored by per-residue confidence (pLDDT). Residues are colored according to AlphaFold confidence: dark blue, pLDDT>90 (very high confidence); light blue, 70–90 (confident); yellow, 50–70 (low confidence); and orange, <50 (very low confidence). High-confidence regions likely correspond to well-defined structural elements, whereas low-confidence regions may represent flexible or intrinsically disordered segments. **B.** Prediction of intrinsically disordered regions (IDRs) in DTH1 using PONDR algorithms. The blue line shows disorder probability profiles from Various Long 3 (VL3), and the purple line from Various Short Long 2 (VSL2) predictions. Most of DTH1 is predicted to be disordered (PONDR score > 0.5). **C.** Schematic representation of the domain architecture of DTH1, with the coiled-coil region shown in blue, which also contains the lipid-binding domain (LBD) previously identified by (Lee *et al*., 2020), and the C-terminal conserved alpha-helices shown in orange. **D.** Alphafold3 prediction of the α-helical domain alone found in the C-terminal of DTH1. The color scheme is the same as in panel A.

The only structured part predicted by AlphaFold3 is an extensive coiled-coil domain (CC-domain) (**Figure 1A**) comprising 800 amino acids (**Figure 1C**), which constitutes nearly half of the protein sequence. The CC-domain encompasses the previously identified LBD domain (247 amino acids). Given the prominence of the CC-domain within DTH1, this structural element is likely central to its function. For example, it may facilitate the recruitment of proteins to the LD surface or mediate interactions with other organelles. Coiled-coil domains are known to promote membrane tethering and fusion events, such as those involved in LD-lysosome/vacuole interactions, analogous to mechanisms observed for viral membrane fusion proteins (Harrison, 2015).

Among other features, at the C-terminus of DTH1, we find an α-helical domain (αH-domain) as predicted by the Jalview JPRED secondary structure prediction tool. Interestingly, this αH-domain also appears in DTH1 homologs (**Supplemental Figure S1**), suggesting a conserved, specific function. To gain further insight into the three-dimensional organization of this region, we analyzed the C-terminal portion of DTH1 using AlphaFold3. The resulting model predicted a structure containing multiple closely packed α-helices arranged into a cage-like architecture (**Figure 1D**). The structural organization could potentially facilitate the trapping or scaffolding of protein partners or alternatively participate in lipid transfer/export processes. Taken together, the above in-silico analyses support the relevance of DTH1 as a promising target for more detailed structural and functional characterization.

### LD lipidomics highlights the role of PE in lipid degradation

To promote LD turnover, both the LD membrane lipid monolayer and the proteins coating or associated with the LD surface should be targeted for degradation (D’Andrea, 2016; Deruyffelaere et al., 2018; Kretzschmar et al., 2018). To further investigate the steps involving DTH1 in LD degradation, we first performed whole-cell lipidomic analysis of WT (CC4533) and the two *dth1* mutants *(dth1-2* and *dth1-3*) during TAG accumulation (TAP-N [NR0]) and degradation phase (MM+N [NR5]) (**Supplemental Figure S2A**). Apart from the difference already observed in TAG degradation (as previously reported in Lee *et al*., 2020), no significant difference in whole-cell membrane lipid classes was found in either condition (**Supplemental Figure S2B**). Looking into lipid molecular species, *dth1* mutants have significantly higher amounts of plastidial lipids under N-recovery, particularly DGDG34:4, MGDG34:4, MGDG34:5, PG34:3, and SQDG34:2 (**Supplemental Figure S2C**). This increase may reflect an altered chloroplast lipid metabolism in *dth1* mutants, or a compensatory response to the impaired fatty acid flux from TAG turnover to rebuild the thylakoid membrane under N resupply conditions.

Since DTH1 is located on LDs, we isolated LDs from both WT and *dth1* mutants from the same batch as for whole-cell lipidomic analysis (48 h TAP-N [NR0] and 5 h MM+N [NR5]) and performed lipidomics analysis (**Figure 2A**). The lipid composition of LD membrane coat has been reported for conditions during active LD biogenesis (TAP-N) (Tsai *et al*., 2015), but has not been investigated during LD remobilization. Here, LC-MS analysis of membrane lipids from isolated LDs during remobilization revealed the presence of all major known LD membrane lipid species, with DGTS being the most abundant lipid of the LD monolayer, followed by DGDG (**Figure 2B**). The comparison of the LD monolayer lipid composition of NR0 and NR5 in the WT revealed a significant increase in the proportion of phosphatidylethanolamine (PE) during LD degradation (**Figure 2B**). Given that DTH1 is a PE-binding protein on LDs (Lee *et al*., 2020), we investigated whether the absence of DTH1 altered this remodeling. During N resupply, PE was significantly less abundant in LDs isolated from both *dth1* mutant lines compared to WT (**Figure 2C**). Looking into detailed PE molecular species, a decrease in each PE species (albeit not significant at the individual level) was noticed in the *dth1* mutants (**Figure 2D**). In addition, three plastidial lipids (DGDG34:4, SQDG34:2 and SQDG34:3) showed higher relative abundance in *dth1* mutants under N resupply. Taken together, these results demonstrate that the LD membrane undergoes a dynamic remodeling during early steps of LD degradation and point to an important role for PE in this process.

**Figure 2.**
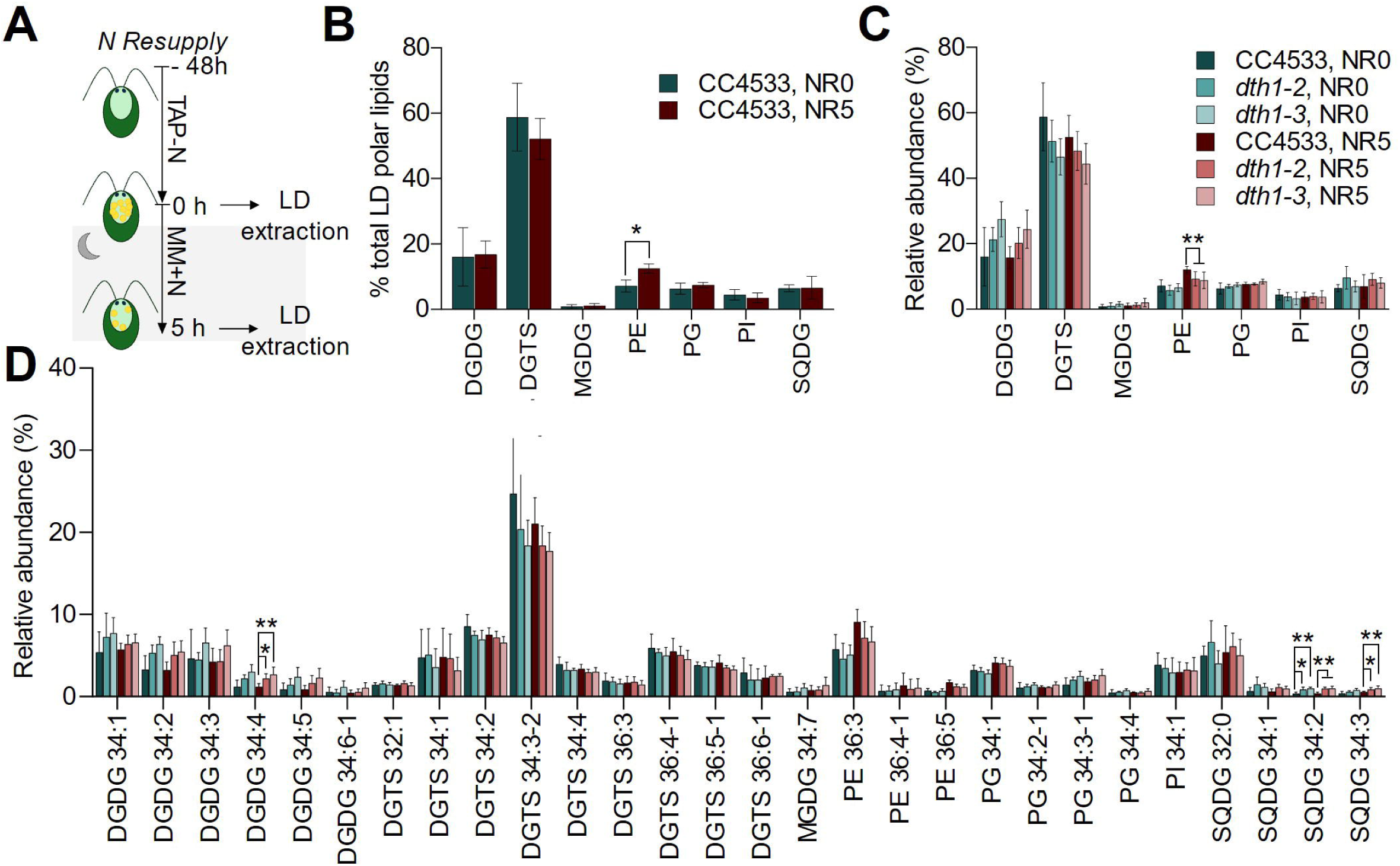
Lipidomics analyses of WT and *dth1* mutants during N starvation and N recovery. **A.** Experimental workflow. LDs were extracted from cells after 48 h of nitrogen starvation (NR0) and after 5 h of nitrogen resupply (NR5). Lipids were separated into neutral (NL) and polar (PL) fractions, with only the polar fraction analyzed. **B.** LC-MS analysis of LD polar lipid composition in wild-type (CC4533) under NR0 and NR5 conditions. Data are presented as mean ± standard deviation (n = 8 biological replicates). Significance was tested using the Wilcoxon rank-sum test, with p-values adjusted for multiple comparisons via the Benjamini–Hochberg false discovery rate (FDR) correction (*\*p* < 0.05). **C.** LC-MS analysis of LD polar lipid class composition in WT and *dth1* mutants under NR0 and NR5 conditions. Lipid MS peak areas were summed by class and normalized against total peak area per replicate to determine the percentage contribution of each lipid class. **D.** LC-MS analysis of LD polar lipid species composition in WT and *dth1* mutants under NR0 and NR5 conditions. Lipid MS peak areas of each species were divided by total peak areas per replicate to calculate the percentage contribution of each lipid species. Species represented by < 0.5% of total response were omitted. Data in panels (C) and (D) are shown as mean ± SD from 6 to 10 biological replicates; significance was assessed using the Kruskal–Wallis test followed by Dunn’s multiple comparison test, with *p-*values adjusted via FDR correction (*\*p* < 0.05; *\*\*p* < 0.01). Abbreviations: DGDG, digalactosyldiacylglycerol; DGTS, diacylglyceryl-*N,N,N*-trimethylhomoserine; MGDG, monogalactosyldiacylglycerol; PE, phosphatidylethanolamine; PG, phosphatidylglycerol; PI, phosphatidylinositol; SQDG, sulfoquinovosyldiacylglycerol.

### LD proteomics identify substantial proteome remodeling during N resupply

Although the LD proteome has been reported several times (Moellering and Benning, 2010; Nguyen *et al*., 2011; Goold *et al*., 2016; Torres-Romero *et al*., 2024), no report is yet available on the LD proteome under active TAG degradation. To fill this gap, we performed mass spectrometry (MS)-based label-free quantitative proteomics on LD-enriched fractions isolated from N-starved (48 h TAP-N, [NR0]) and N-resupplied (12 h MM+N, [NR12]) wild-type (CC4533) and *dth1* mutant cells, using five biological replicates per condition. We specifically chose this time point because the number of LDs remained sufficiently abundant for purification after 12 h of N resupply (**Figure 3A-B**).

**Figure 3.**
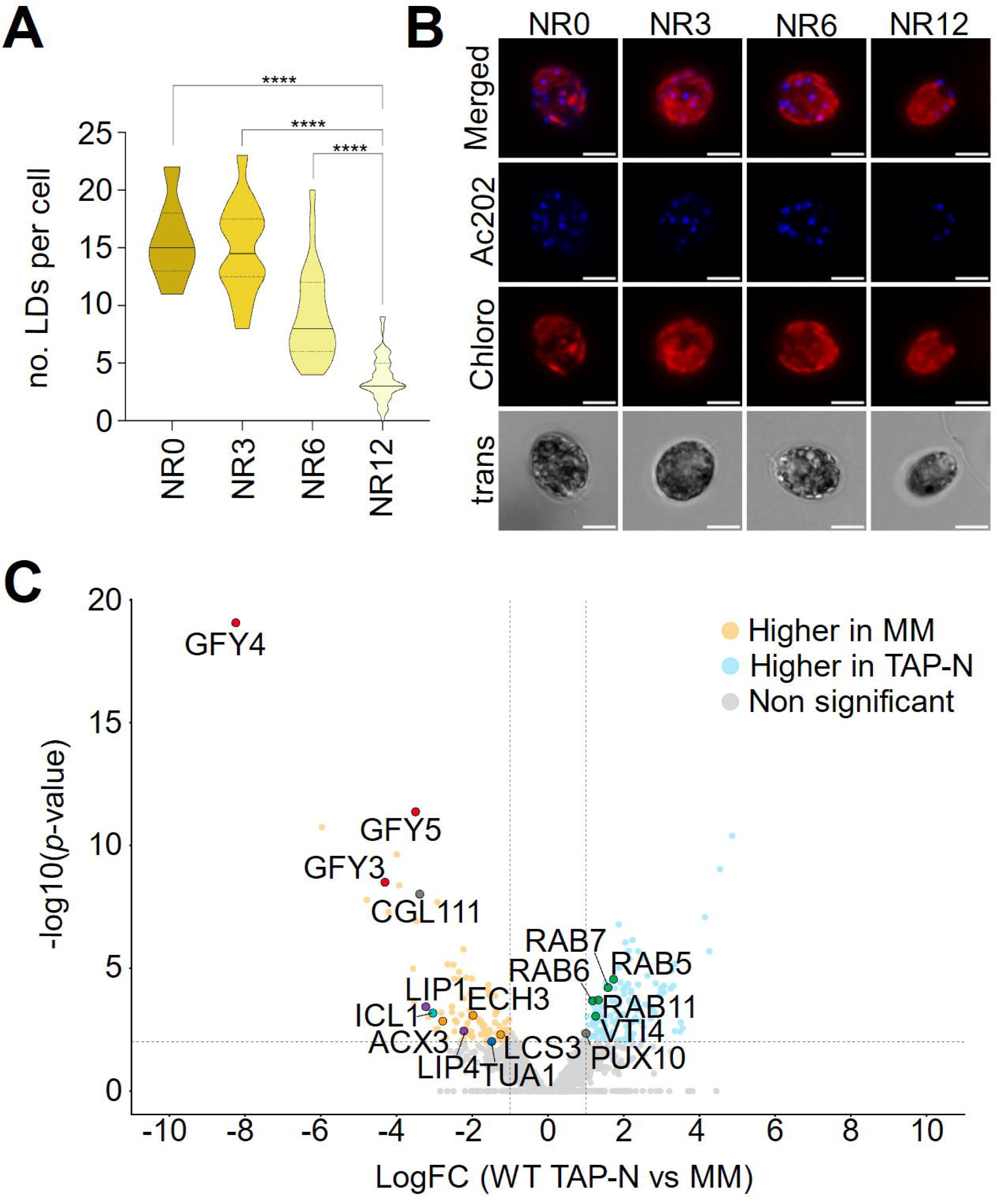
MS-based quantitative proteomic analysis of LDs purified from wild-type (CC4533) Chlamydomonas cells during N starvation and N resupply conditions. **A.** Quantification of LD sizes in WT during N starvation and N resupply conditions. Areas of AC202-stained LDs were measured using ImageJ (refer to “Materials and methods” for details), and values shown in violin plots represent those obtained from n=7-50 independent cells from the same biological replicate, including those shown in (C). Statistical analysis was assessed by Kruskal–Wallis and Dunn’s test with FDR correction (*\*\*\*\*p* < 0.0001). **B.** Representative confocal images of WT under N starvation (NR0) and N resupply (NR3, NR6 and NR12). LDs were stained with AC202 (blue) and chlorophyll autofluorescence is shown in red. Bars = 5 µm. **C.** Volcano plot displaying the differential abundance of proteins analyzed by MS-based label-free quantitative proteomics (n=5 per condition). The volcano plot represents the -log10 (limma *p*-value) on the y axis plotted against the log2(Fold Change NR12 vs NR0) on the x-axis for each quantified protein. Blue and yellow dots represent proteins found significantly enriched in NR0 and NR12 conditions, respectively (log2(Fold Change) ≥ |1| and -log10(p-value) ≥ 2, leading to a Benjamini-Hochberg FDR = 2.27%). Proteins involved in LD homeostasis are labeled and grouped into different classes: grey, protein degradation; green, vesicular trafficking; blue, cytoskeleton; orange, fatty acid beta-oxidation pathway; red, transporters; purple, lipases and turquoise, glyoxylate cycle.

The proteomic analyses allowed to reliably identify and quantify 1559 proteins across the 30 LD samples (6 genotypes/condition with 5 biological replicates for each genotype) (**Supplemental Table S1A**). Among the most abundant proteins detected in both conditions (**Supplemental Table S1B**), there were several well characterized LD-associated proteins, including the MLDP, the betaine lipid synthase (BTA1), the lyso-diacylglyceryltrimethylhomo-Ser (lyso-DGTS) lipase (ABHD1), the glycosyl hydrolase (GHL1), the lysophospholipid acyltransferase PGA2 (also known as LPLAT2) and the long-chain acyl-CoA synthetase (LCS2), consistent with previous reports (Moellering and Benning, 2010; Nguyen *et al*., 2011; Goold *et al*., 2016; Torres-Romero *et al*., 2024).

The differential abundance analysis between LDs isolated under TAP-N (NR0) and MM+N (NR12) conditions (log₂FC > |1|, limma *p* < 0.01, Benjamini-Hochberg FDR = 2.27%) identified 91 proteins significantly enriched and 118 proteins significantly depleted under N resupply **(Figure 3C; Supplemental Table S1A**). Among the proteins enriched under N-resupply compared to N-starvation condition, we notably identified a phosphatidylcholine (PC) transfer protein (CGL111 [Cre03.g174850]) that could be involved in removing DGTS (a PC analog in Chlamydomonas) from the LD monolayer. Additionally, a putative phospholipase with a DUF676 domain (Cre05.g233850) was found, homologous to the phospholipase B LPL1 identified as playing a role in LD membrane coat degradation in the budding yeast (Selvaraju *et al*., 2014). We also detected several enzymes known or presumed to be involved in lipid catabolism. These include the lipases LIP4 and LIP1 (Li *et al*., 2012; Warakanont *et al*., 2019), LCS2 (Bai *et al*., 2022), the acyl-CoA oxidase 3 (ACX3) and the 3-hydroxyacyl-CoA dehydrogenase (ECH3) (Kong *et al*., 2017), among other lipid-associated proteins. In addition, we identified two key components of the glyoxylate cycle, the isocitrate lyase 1 (ICL1) and the aconitate hydratase 1 (ACH1) (Plancke *et al*., 2014; Lauersen *et al*., 2016), as well as the gluconeogeneic enzyme transaldolase (TAL1) (Goodenough *et al*., 2014). These findings suggest a coordinated recruitment of lipid catabolic and metabolic reprogramming enzymes to LD-associated fractions during N resupply.

Interestingly, among the most strongly enriched proteins under MM+N (NR12) condition compared to N starvation, we identified three putative acetate transporters belonging to the GPR1/FUN34/YaaH family (GFY4, 3, 5) (**Figure 3C**). Their role in acetate transport has not yet been conclusively demonstrated (Durante *et al*., 2019), and more intriguingly the abundance of these proteins remains similar regardless of the acetate levels in the media (Füßl *et al*., 2022). In contrast, their protein abundance is markedly increased upon transfer from light to dark, a condition associated with enhanced catabolic activity (Füßl *et al*., 2022). Collectively, these observations suggest that GFY3-5 may play a role in intracellular carbon trafficking rather than in acetate uptake from the media.

Interestingly, the relative abundance of the major structural LD protein MLDP decreased from ∼23% to ∼15% of total LD-associated proteins (iBAQ values) when comparing LDs isolated from N-starved cells with those from N-resupplied WT cells. This observation is consistent with the idea that major structural LD proteins, such as oleosin or MLDP, are selectively removed during active LD degradation, thereby increasing lipolytic enzyme access to the TAG core (Tsai *et al*., 2015; Deruyffelaere *et al*., 2018; Kretzschmar *et al*., 2018). In addition, alpha-tubulin TUA1, previously reported to interact with MLDP (Tsai et al., 2015), was more abundant during active LD degradation, consistent with a possible requirement for increased LD mobility, possibly toward peroxisomes or vacuoles (Thazar-Poulot *et al*., 2015; Singh *et al*., 2022). Supporting this, we identified a VAMP-associated protein (Cre03.g196900) among the more enriched proteins during N-resupply, which is homologous to VAP27 in *Arabidopsis thaliana*. VAP27 is involved in multiple organelle contact sites, notably between the endoplasmic reticulum (ER) and mitochondria, where it interacts with the mitochondrial protein TRB1 to mediate the selective autophagy of damaged mitochondria (Li *et al*., 2022). This process might be extended to include lipophagy, i.e., LD-selective autophagy.

Among the proteins more abundant under TAP-N (NR0) condition, the most strongly enriched in the TAP-N versus MM+N (NR12) condition was a putative wax synthase belonging to the membrane-bound O-acyl transferase (MBOAT) (**Supplemental Table S1A**). Neither its function in Chlamydomonas nor its homologs in other species have been reported. In addition, we detected a couple of nitrogen/nitrate transporters, multiple Rab GTPases (RAB5, RAB6, RAB7, RAB11 and RAB1-like), and one Q-SNARE protein (VTI4) (**Figure 3C**), consistent with vesicular trafficking activity during LD formation, likely regulating fusion, fission and membrane contact with other organelles. Interestingly, we also identified a protein homologous to PUX10 (**Figure 3C**), a member of the plant UBX domain-containing (PUX) protein family previously described in *Arabidopsis thaliana* for its role in LD degradation (Kretzschmar *et al*., 2018). Taken together, the LD proteome of Chlamydomonas undergoes extensive remodeling during N-resupply. The upregulation of enzymes of lipid catabolism and GFY proteins suggests an increased need for active lipid degradation and intracellular carbon trafficking during stress recovery.

### The absence of DTH1 induces significant changes in the LD proteome

We further explored the role of DTH1 in LD degradation, comparing proteomes of LDs extracted from CC4533 and *dth1* mutants during N starvation (48 h TAP-N, [NR0]) and N resupply (12 h MM+N, [NR12]) conditions. Considering proteins found differentially abundant during LD formation (NR0) in WT compared to both *dth1* mutants, we found 5 proteins significantly enriched in LDs purified from WT and 6 proteins enriched in LDs purified from both *dth1* mutants (**Figure 4B**; **Supplemental Table S1A**; **Table 1**). The five proteins more abundant in CC4533 include the GFY5 (Durante *et al*., 2019), a mitochondrial fatty acid anion carrier protein (MCP28/UCP2C), PUX10, one mitochondrial protein with unknown function and finally a peroxisomal biogenesis factor 11 (PEX11). Interestingly, all above five proteins are known or presumably to be proteins of either mitochondrial or peroxisomal origin, suggesting that DTH1 might facilitate in bringing these organelles closer to LDs. In the absence of DTH1 (i.e. *dth1* mutants), proteins found enriched compared to CC4533 are a flagellar associated protein (FAP325), a xanthine/uracil/vitamin C permease-like (XUV2), a flagellar membrane glycoprotein (FMG2), a chloroplast fatty acid desaturase (FAD6), a membrane-associated progesterone receptor component-related protein, and an unknown protein (Cre18.g748297). The latter shows ∼30% similarity to another protein (Cre09.g404750) annotated as a scavenger receptor cysteine-rich (SRCR) protein. Taken together, most of proteins found enriched in CC4533 during LD formation were mitochondrial or peroxisomal, suggesting that the presence of DTH1 on LD surface favors contact with these organelles during active LD formation.

**Figure 4.**
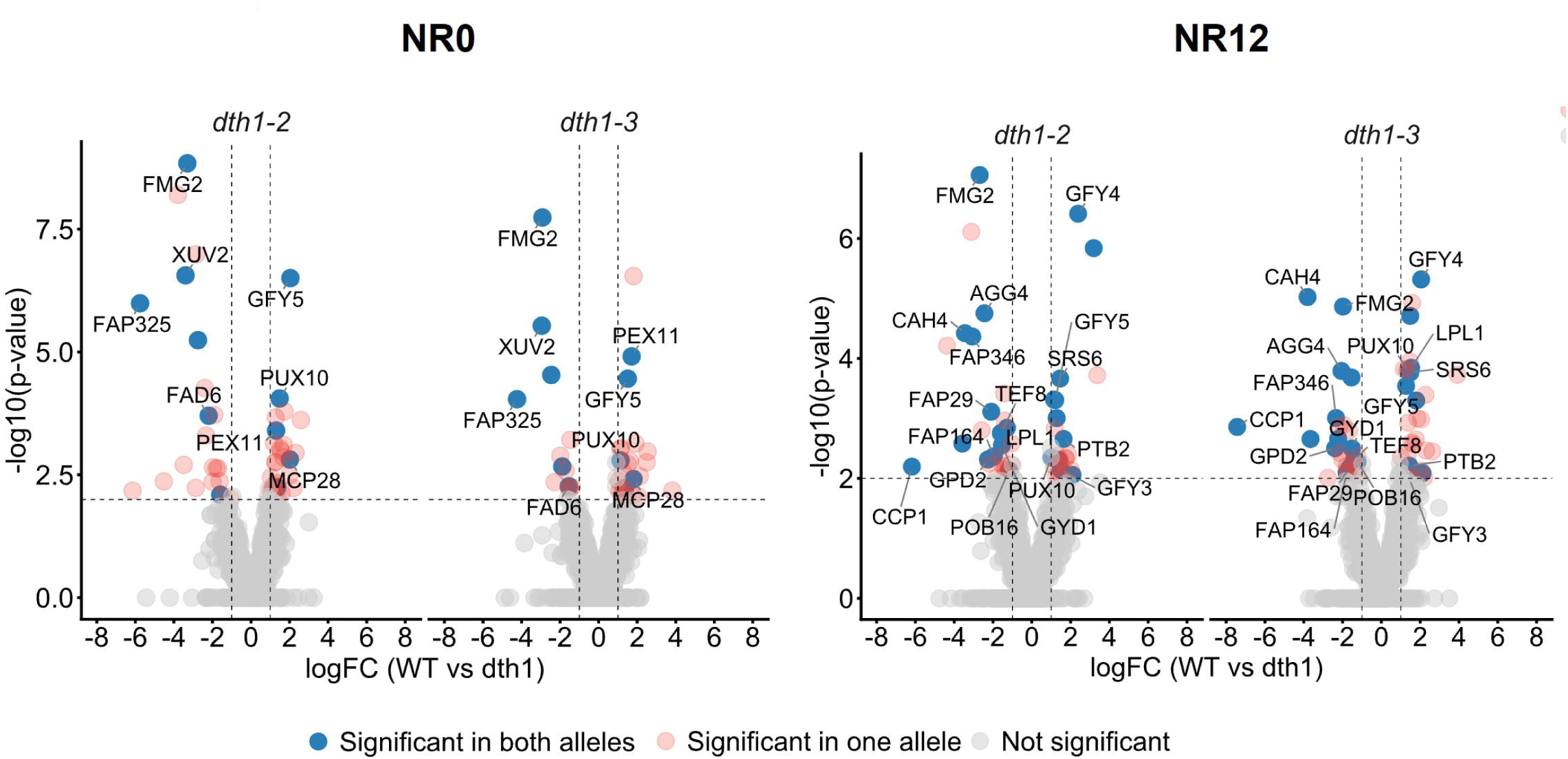
MS-based quantitative proteomics of LDs purified from WT and *dth1* mutants during N starvation and N resupply. Volcano plots of the differential abundance of proteins identified in CC4533 versus *dth1* mutants under N starvation (NR0) and N resupply (NR12) conditions. Protein identification and abundance were determined by MS-based label-free quantitative proteomics (n = 5 per condition). The volcano plots display -log10(limma p-value) on the y-axis versus log2(fold change) on the x-axis for each quantified protein. Dark blue points denote proteins significantly enriched in both mutants (log2(fold change) ≥ |1| and -log10(p-value) ≥ 2, corresponding to Benjamini-Hochberg FDRs inferior to 5.3 %); light red points denote proteins significant in only one mutant; grey points denote non-significant proteins. Vertical dashed lines indicate log2(fold change) ≥ |1|, and the horizontal dashed line indicates – log10(p-value) ≤ 2 (p-value ≤ 0.01).

**Table 1:** Significantly differentiated proteins in WT versus *dth1* mutants during N starvation (NR0).

|  | ID | Gene name | Description | CC4533 NR0 vs *dth1-2* |  | CC4533 NR0 vs *dth1-3* |  |
| --- | --- | --- | --- | --- | --- | --- | --- |
|  |  |  |  | logFC | p-value | logFC | p-value |
| +++ in *dth1* | Cre06.g263200 | FAP325 | Flagellar Associated Protein 325 | -5.76 | 1.02E-06 | 4.23 | 9.14E-05 |
|  | Cre10.g448450 | XUV2,UAPA5 | Xanthine/uracil/vitamin C permease-like | -3.40 | 2.75E-07 | -2.95 | 2.92E-06 |
|  | Cre09.g392867 | FMG2 | Flagella Membrane Glycoprotein 1B | -3.30 | 1.44E-09 | -2.92 | 1.83E-08 |
|  | Cre18.g748297 |  | No clear annotation | -2.75 | 5.74E-06 | -2.46 | 2.93E-05 |
|  | Cre13.g590500 | FAD6 | Omega-6-fatty acid desaturase, chloroplast isoform | -2.19 | 1.98E-04 | -1.55 | 5.42E-03 |
|  | Cre03.g148100 |  | Membrane-associated progesterone receptor component-related protein | -1.59 | 8.01E-03 | -1.89 | 2.14E-03 |
| +++ in WT | Cre06.g263300 | PEX11 | Peroxisomal biogenesis factor 11 | 1.30 | 3.97E-04 | 1.71 | 1.23E-05 |
|  | Cre12.g550450 | Neuferricin | Membrane-associated progesterone receptor component-related protein | 1.36 | 5.70E-03 | 1.35 | 5.97E-03 |
|  | Cre03.g200100 | PUX10 | Plant UBX domain-containing protein 10 | 1.49 | 8.83E-05 | 1.14 | 1.14 |
|  | Cre15.g641200 | MCP28 | Mitochondrial substrate carrier protein | 2.01 | 1.54E-03 | 1.81 | 3.87E-03 |
|  | Cre17.g702950 | GFY5 | GPR1/FUN34/YaaH family membrane protein | 2.04 | 3.13E-07 | 1.51 | 3.48E-05 |

During LD degradation (NR12), 11 proteins were found significantly enriched in LDs purified from CC4533 while 14 proteins were found more abundant in LDs purified from both *dth1* mutants (**Figure 4D; Supplemental Table S1A; Table 2**). Proteins more abundant in LDs obtained from CC4533 included the three GFY(3, 4, 5) proteins as well as two homologs of known LD-associated proteins: PUX10 and the phospholipase LPL1-like, homolog to the phospholipase B LPL1 in yeast (Selvaraju *et al*., 2014; Weisshaar *et al*., 2017). Interestingly, LIP4 was significantly and strongly enriched in WT compared with *dth1-2*. A similar enrichment was observed in WT relative to *dth1-3*, although the *p*-value was just below our significance threshold (*p* = 0.014). Given that *dth1* mutants are strongly impacted in LD remobilization, this result remains noteworthy, as LIP4 is a TAG lipase involved in LD degradation (Warakanont *et al*., 2019). Surprisingly, several proteins associated with the low-CO_2_ response in the mitochondria (i.e., the chloroplast carrier protein 1 (CCP1), carbonic anhydrase 4 (CAH4) and glycolate dehydrogenase 1 (GYD1) (Dao *et al*., 2025) were enriched in LDs purified from the *dth1* mutants (**Supplemental Table S1A, Table 2**). Because *dth1* mutants were blocked in TAG catabolism, and therefore presumably reduced CO_2_ production through lipid respiration, these cells may experience an intracellular low-carbon state that triggers the induction of low-CO_2_ responsive proteins. It is worth noting that many of these proteins are peroxisomal or mitochondrial, suggesting a close physical and metabolic association between LDs, peroxisomes and mitochondria during LD degradation. Since peroxisome is the primary site of fatty acid β-oxidation (Kong *et al*., 2017) and the glyoxylate cycle in green algae (Plancke *et al*., 2014; Lauersen *et al*., 2016), the proximity of LDs to these organelles point to a tight metabolic coupling during lipid mobilization and carbon recycling. Indeed, mitochondria and peroxisomes have been seen to be located in proximity to LDs during N resupply (Goodenough and Engel, 2023).

**Table 2:** Significantly differentiated proteins in WT versus *dth1* mutants during LD degradation (NR12).

**degradation (NR12).**
|  | ID | Gene name | Description | CC4533 NR12 vs *dth1-2* |  | CC4533 NR12 vs *dth1-3* |  |
| --- | --- | --- | --- | --- | --- | --- | --- |
|  |  |  |  | logFC | P_Value | logFC | P_Value |
| +++ in *dth1* | Cre04.g223300 | CCP1 | Low-CO<sub>2</sub>-inducible chloroplast envelope protein | -6.18 | 6.37E-03 | -7.43 | 1.39E-03 |
|  | Cre11.g467531 |  |  | -3.58 | 2.63E-03 | -3.65 | 2.21E-03 |
|  | Cre05.g248400 | CAH4 | Mitochondrial carbonic anhydrase, beta type | -3.45 | 3.79E-05 | -3.80 | 9.47E-06 |
|  | Cre12.g492600 | FAP346 | FAS Domain Flagellar Associated Protein 346 | -3.07 | 4.32E-05 | -2.34 | 9.84E-04 |
|  | Cre09.g392867 | FMG2 | Flagella Membrane Glycoprotein 1B | -2.69 | 8.75E-08 | -1.98 | 1.37E-05 |
|  | Cre17.g738050 | AGG4 | Aggregation 4 | -2.44 | 1.76E-05 | -2.06 | 1.62E-04 |
|  | Cre01.g053000 | GPD2 | Glycerol-3-phosphate dehydrogenase | -2.26 | 4.85E-03 | -2.39 | 3.14E-03 |
|  | Cre08.g364300 | FAP29 | Flagellar Associated Protein 29 | -2.09 | 7.74E-04 | -1.63 | 6.60E-03 |
|  | Cre13.g575701 |  | cold regulated protein? | -2.04 | 4.47E-03 | -2.22 | 2.19E-03 |
|  | Cre17.g735350 | FAP164 | Ankyrin Repeat Flagellar Associated Protein 164 | -1.96 | 4.29E-03 | -1.79 | 8.14E-03 |
|  | Cre03.g182150 | TEF8 | predicted protein | -1.60 | 1.75E-03 | -1.41 | 4.95E-03 |
|  | Cre06.g288700 | GYD1 | Glycolate dehydrogenase | -1.54 | 2.81E-03 | -1.52 | 3.16E-03 |
|  | Cre11.g467759 |  | Similar to AtTic21/PIC1 | -1.28 | 1.43E-03 | -1.54 | 2.08E-04 |
|  | Cre17.g696850 | POB16 | Proteome of basal body 16 | -1.17 | 7.31E-03 | -1.23 | 5.30E-03 |
| +++ in WT | Cre03.g200100 | PUX10 | Plant UBX domain-containing protein 10 | 1.01 | 4.50E-03 | 1.41 | 1.64E-04 |
|  | Cre12.g527650 |  |  | 1.14 | 4.85E-04 | 1.49 | 1.95E-05 |
|  | Cre17.g702950 | GFY5 | GPR1/FUN34/YaaH family membrane protein | 1.21 | 4.94E-04 | 1.27 | 2.87E-04 |
|  | Cre05.g233850 | LPL1 | DUF676 DOMAIN-CONTAINING PROTEIN | 1.28 | 9.88E-04 | 1.53 | 1.43E-04 |
|  | Cre05.g240533 |  | E3 ubiquitine ligase | 1.39 | 4.93E-03 | 1.79 | 5.02E-04 |
|  | Cre13.g586916 | SRS6 | Serine/arginine-rich pre-mRNA splicing factor | 1.46 | 2.16E-04 | 1.49 | 1.72E-04 |
|  | Cre17.g700750 | GFY3 | GPR1/FUN34/YaaH family membrane protein | 1.48 | 7.17E-03 | 1.42 | 9.56E-03 |
|  | Cre07.g325741 | PTB2 | Sodium/phosphate symporter | 1.64 | 2.20E-03 | 1.44 | 6.14E-03 |
|  | Cre12.g548200 |  | Peroxisomal leader peptide-processing protease | 2.10 | 8.75E-03 | 2.12 | 8.18E-03 |
|  | Cre17.g702900 | GFY4 | GPR1/FUN34/YaaH family membrane protein | 2.39 | 3.87E-07 | 2.05 | 4.81E-06 |
|  | Cre09.g388726 |  | HAD like hydrolase | 3.19 | 1.45E-06 | 1.52 | 7.41E-03 |

### N resupply induces distinct subcellular changes in both WT and *dth1* mutants

We previously showed that *dth1* mutants contain a higher number of LDs than WT cells during recovery from N starvation (Lee *et al*., 2020), but the accompanying changes in subcellular structures have not been examined. To investigate the subcellular changes associated with DTH1 deficiency, we performed both confocal and transmission electron microscopy (TEM). We quantified both the size and number of LDs per cell during N fluctuation. During N starvation, we observed no significant difference in LD number and size between WT and *dth1* mutants. However, during N resupply, *dth1* mutants retained significantly more and larger LDs than CC4533 (**Figure 5A-B**), highlighting a delay in LD remobilization following N starvation.

**Figure 5.**
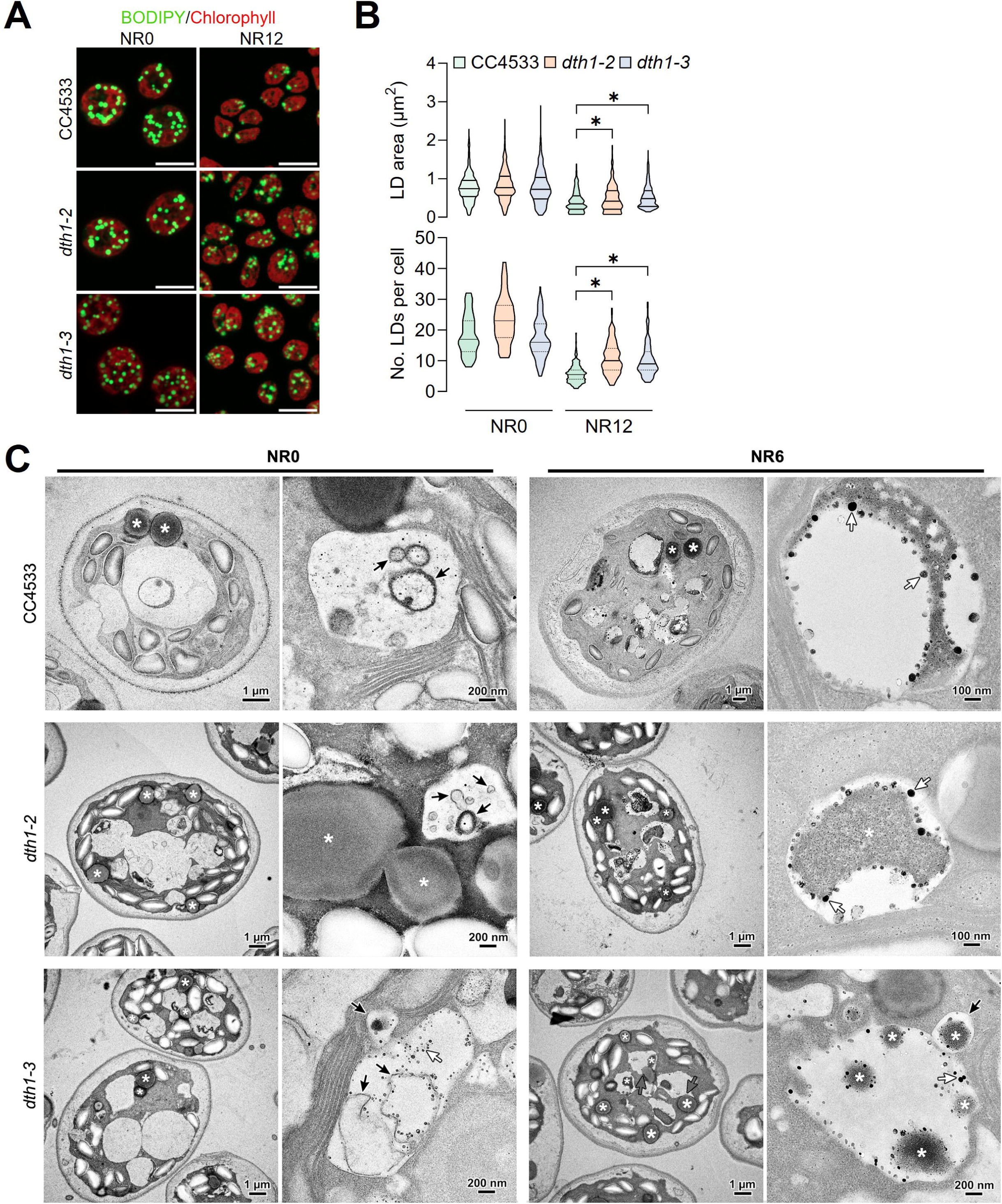
Ultrastructural analysis of WT and *dth1* mutants under N starvation and N resupply conditions. **A.** Confocal images of CC4533, *dth1-2* and *dth1-3* cells under N starvation (NR0) and N resupply (NR12) conditions. Lipid droplets were stained with Bodipy and autofluorescence of the chlorophyll is shown in red. scale bar = 10 µm. **B.** Violin plot of LD size and number distribution across CC-4533 and *dth1* mutants under N starvation (NR0) and N resupply conditions (NR12). Data represent n=47 individual cells (CC4533, NR0), n=33 (*dth1-2*, NR0), n=115 (*dth1-3*, NR0), n=170 (CC4533, NR12), n=157 (*dth1-2*, NR12) and n=134 (*dth1-3*, NR12). Statistical analyses were conducted using a Kruskal–Wallis test followed by Dunn’s multiple comparison test, with p-values adjusted via FDR correction (*\*p* < 0.05). **C.** TEM images of CC4533, *dth1-2* and *dth1-3* under NR0 and NR6 conditions. LDs are electron-dense spherical structures (white asterix) and vacuoles are the clear cytoplasmic regions. The red arrow in *dth1-3*, NR6 condition represents large LDs integrating vacuoles through a process resembling macrolipophagy; orange arrows represent micro-LDs (mLDs) and blue arrows point to autophagosome-like structures.

To examine subcellular structures in greater detail, we used TEM to visualize LD degradation following N resupply. TEM provides high-resolution images of the intracellular organization of organelles (Tsai *et al*., 2015). We have examined two time points: at the height of N starvation (NR0) and during N-resupply (NR6). After a quick spin, cells were immediately fixed in glutaraldehyde (2.5%) to preserve cellular ultrastructure for TEM analysis. N starvation (NR0) led to the accumulation of starch and LDs and the formation of large vacuoles, often central and occupying nearly the entire cytoplasm (Moellering and Benning, 2010; Tsai *et al*., 2018) (**Figure 5C**). Those vacuoles are filled with structures resembling autophagosomes (**Figure 5C**, black arrows NR0), sometimes containing small LDs (**Figure 5C**, white asterisk). Vacuolar lytic activity is required for TAG synthesis and LD formation in N-starved cells (Couso *et al*., 2018), which could explain this observation. During recovery from N starvation (NR6), we observed increased cytoplasmic vacuolization in both WT and *dth1* mutants, characterized by an increase in the number of smaller lytic vacuoles (LV) (**Figure 5C**). Most of these contained LDs undergoing degradation (**Figure 5C**), which appear to enter the vacuole either via encapsulation of small LDs within autophagosomes followed by fusion with the LV (i.e., macrolipophagy, **Figure 5C**, black arrow in *dth1-3* NR6), or through direct fusion of larger LDs with the vacuole (i.e., macrolipophagy, **Figure 5C**, grey arrows in *dth1-3* NR6). Consistent with the confocal images, we observed an increased number of intact LDs inside *dth1* mutants compared to WT cells after 6 h of N-resupply (**Figure 5C**).

In both conditions, we observed the appearance of small LDs, referred to in the literature as micro-LDs (mLDs), in all strains (**Figure 5C**, white arrows). Notably, mLDs were predominantly detected in LV from both WT and *dth1* cells, arguing against a role for DTH1 in LD delivery or tethering to the vacuole. The formation of mLDs has previously been reported in mammalian cells under lipolytic conditions (Ariotti *et al*., 2012). Such LD remodeling may increase the LD surface-to-volume ratio, thereby improving lipase accessibility and enhancing TAG hydrolysis (D’Andrea, 2016). The formation of mLDs may result either from the fragmentation of larger LDs (Moore *et al*., 2005) or from the reformation of small LDs during TAG degradation to buffer the release of potentially toxic free fatty acids (FFAs) (Paar *et al*., 2012). Moreover, in both WT and *dth1* mutants, we observed that LDs move toward the periplasmic space (**Supplemental Figure S3A-C**). Concomitant with this, we observed mLDs in the periplasmic space as well (**Figure S3D-E**), which can either come from the LD degradation in the periplasmic space (**Figure S3A, F**) or from the exocytosis of vesicular materials (**Figure S3G**). We also observed those electron-dense structures around starch granules (**Figure S3H**) and in flagella (**Figure S3I**) under N resupply conditions. To our knowledge, this is the first report of mLD formation in algal cells. Collectively, these results indicate that N resupply triggers distinct subcellular remodeling in both WT and *dth1* cells, following a similar pattern between the strains, except for a higher number of LDs observed in *dth1* mutants under N resupply conditions.

## Discussion

All living organisms have evolved ways to cope with environmental challenges including both abiotic and biotic stresses. Among the various adaptive responses, lipid homeostasis encompassing membrane lipid remodeling and LD homeostasis plays a central role (Wang *et al*., 2019; Chapman *et al*., 2012; Ischebeck *et al*., 2020). Here, we showed that the lipidome and proteome of LDs are under active remodeling during nutrient fluctuation. We identified PE as an important lipid during LD degradation and showed that the LD proteome undergoes extensive remodeling during recovery from N starvation, including enrichment in lipid-degradation enzymes, and in proteins of mitochondrial and peroxisomal origins to ensure proper intracellular carbon usage.

### Proteins of lipolysis, fatty acid β-oxidation and glyoxylate cycle are enriched in LD preparations under active degradation

The functions of LDs are largely mediated by their associated proteins. While some proteins are constitutively associated with the LD surface, others are dynamically recruited in response to changing cellular conditions through protein-protein or protein-lipid interactions (Dhiman *et al*., 2020). We found that the Chlamydomonas LD proteome undergoes dynamic changes following N resupply after a period of N starvation, enhancing the presence of proteins related to lipid catabolism (**Figure 6**). Conceptually, complete LD degradation requires sequential action by distinct sets of proteins. First, to access the carbon and energy stored within LDs, cells must first remodel or dismantle the protein and lipid monolayer surrounding the LD. Related to this first step, we identified a putative phospholipase similar to the phospholipase B LPL1 found in budding yeast (Selvaraju *et al*., 2014), and a homolog of a putative phosphatidylcholine (PC) transfer protein that may remove DGTS (a PC analog in Chlamydomonas) from the LD monolayer. These proteins could destabilize the LD coat, creating space for lipases to access the neutral lipid core. In addition, we identified two previously characterized LD lipases, LIP4 and LIP1 (Li *et al*., 2012; Warakanont *et al*., 2019), enriched on the LD surface during LD degradation.

**Figure 6.**
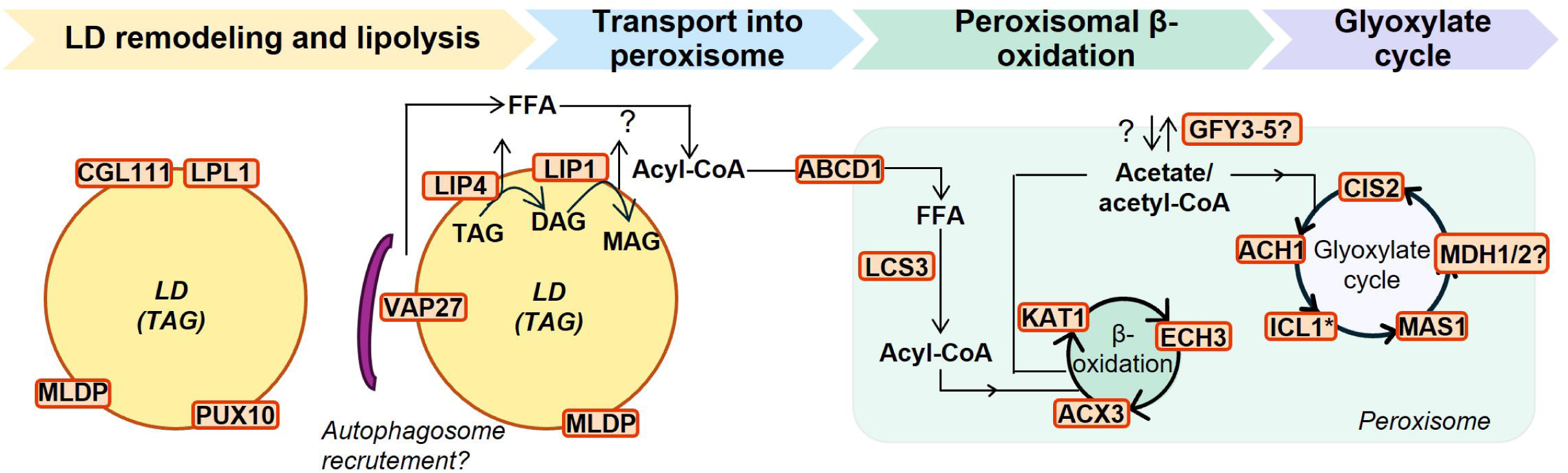
From LD remodeling to peroxisomal β-oxidation: insights from comparative proteomics. The first step is the removal or remodeling of the LD protein and lipid coat to allow lipases access to the LD core, which could be mediated in part by CGL111, a putative DGTS (a PC analog in Chlamydomonas) transferase, and by LPL1, a putative phospholipase B. The next and more critical step is TAG breakdown by lipases: we identified LIP4, a TAG lipase, and LIP1, a DAG lipase, on the LD surface under N resupply conditions. An alternative route for LD degradation is lipophagy (i.e., the selective autophagy of LDs), which requires recruitment of an autophagosome to the LD surface, a process that could be initiated by VAP27, which we found on the LD surface during degradation. Once fatty acids are released from the glycerol backbone, they can either be used for membrane synthesis or taken up by the peroxisome via the ABCD1 for complete β-oxidation. Two β-oxidation enzymes were identified here: ACX3 and ECH3. The resulting acetyl-CoA then enters the tricarboxylic acid cycle in the mitochondria for CO_2_ and ATP production, or enters the glyoxylate cycle to fuel gluconeogenesis. ICL1, a key enzyme of the glyoxylate cycle, was found to be more abundant during N resupply. Finally, putative GFY3,4,5 could act as acetate transporters for shuttling acetate into or out of the peroxisome. Background colors in protein boxes reflects logFC values, ranging from 9 (more abundant in TAP-N, blue) to −9 (more abundant in MM+N, yellow); grey background denotes non-significant proteins but known to be involved in lipid catabolism.

Free fatty acids (FFAs) released by lipolysis can then be used either for building membranes or being completely oxidized through peroxisomal β-oxidation and glyoxylate cycle (Tsai *et al*., 2014, 2015; Kong *et al*., 2017, 2018; Li-Beisson *et al*., 2021). Consistent with this, we found two enzymes of the β-oxidation pathway, ACX3 and ECH3, and one protein of the glyoxylate cycle ICL1, in the LD-enriched fractions during LD degradation. The resulting carbon intermediates are then transferred to mitochondria to fuel ATP production (Tsai *et al*., 2014, 2015; Kong *et al*., 2017, 2018; Li-Beisson *et al*., 2021). In this comparative proteomic analysis, we also identified the enrichment of putative acetate transporters of the GFY family (GFY3-5). Altogether, these results support extensive intracellular carbon trafficking during LD degradation to facilitate coordinated carbon remobilization from lipid to acetyl-CoA to ATP production by transporting carbon intermediates from LDs to peroxisomes and finally to mitochondria for respiration. This is further supported by the enrichment of a number of low CO_2_ response mitochondrial proteins in the TAG degradation-impaired *dth1* mutants. The accumulation of these proteins strongly suggests active trafficking of carbon intermediates from peroxisomes to mitochondria, where they are further metabolized through respiratory pathways and contribute to internal CO_2_ production.

It is important to acknowledge that any organelle purification approaches are inherently susceptible to contamination, but their reproducible co-purification across five biological replicates strongly suggests a strong association (in the presence of 9 M urea), and an enrichment within the immediate LD microenvironment. In particular, the recovery of mitochondrial and peroxisomal proteins is consistent with the existence of close physical contacts and coordinated metabolic interactions between these organelles and LDs during stress recovery (Goodenough and Engel, 2023).

### The importance of PE in LD degradation

During LD transition from accumulation to degradation, relatively higher amount of PE was observed in wild-type Chlamydomonas cells (**Figure 2B**). Interestingly, this enrichment of PE in the LD monolayer is severely impacted by the absence of DTH1. In Chlamydomonas, DTH1 has been demonstrated to be a PE-binding protein and its binding is critical to LD degradation (Lee *et al*., 2020). Our findings suggest that DTH1 recruitment at the LD surface may contribute to PE enrichment during this process. Reduced PE at the LD surface would also limit the anchorage of other PE-dependent actors of LD degradation, such as LC3/ATG8, an autophagy-related protein and a central marker of autophagosome formation (Singh *et al*., 2009). Lipidation of LC3/ATG8 can directly happen on large LDs, where ATG3 lipidates LC3 on LDs after an extended period of starvation, i.e, during the LD degradation process (Omrane *et al*., 2023). A reduction of the PE pool in *dth1* mutant would be expected to reduce LC3/ATG8 lipidation at the LD surface and impair LD degradation through lipophagy. Consistent with PE enrichment acting as an upstream trigger of LD degradation, raising the PE/PC ratio specifically at the LD monolayer is sufficient to activate lipophagy (Jung *et al*., 2023). The small molecule SB2301 translocates PCYT2 to LDs, increases the PE/PC ratio of the LD membrane, and thereby drives LD coalescence and ubiquitin-mediated lipophagy (Jung *et al*., 2023).

Taken together, these observations point collectively to the importance of PE and PE-binding proteins in LD turnover across diverse lineages.

### mLDs as a conserved strategy to buffer excess free fatty acid

Our study provides, to our knowledge, the first evidence for the formation of mLDs in microalgae. The LD size is generally subject to strict cellular control, as it directly influences the surface-to-volume ratio of the organelle and, consequently, the efficiency of lipid remobilization. The generation of mLDs may increase the available surface area for the recruitment and activity of lipases, thus enhancing access to TAG substrates and promoting lipid catabolism. The formation of mLDs during lipoplysis has previously been reported in mammalian cells (Paar *et al*., 2012). Similar structures have also been observed in the leaves of plant mutants impaired in fatty acid import into peroxisomes, where excess cytosolic free fatty acids are transiently re-esterified and stored in newly formed LDs (Park *et al*., 2013). The occurrence of mLDs during N recovery in Chlamydomonas therefore suggests that LD fragmentation or de novo formation of small LDs may represent a conserved strategy to facilitate lipid turnover and maintain lipid homeostasis during periods of intense carbon remobilization.

In conclusion, our study provides a rich dataset on the proteomic and lipidomic remodeling of LDs during their degradation phase in Chlamydomonas. This phase is critical for cell viability, as it ensures the timely remobilization of stored carbon and energy reserves to support the resumption of growth. By capturing the molecular dynamics underlying LD breakdown, our findings shed new light on the mechanisms governing this crucial step of the lipid metabolic cycle. Beyond their fundamental relevance, these results lay the groundwork for the rational design of genetically engineered strains with enhanced oil content, offering promising avenues for the industrial exploitation of Chlamydomonas as a biofuel and bioproduct platform.

## Materials and methods

### Bioinformatics analyses

The DTH1 protein sequence (from transcript 2) was retrieved from the JGI Phytozome 14 database (https://phytozome-next.jgi.doe.gov/). Basic sequence and structural features, including disordered regions, coiled-coils, and potential signal peptides, were predicted using multiple software tools: PredictProtein (https://predictprotein.org/), Dismeta (https://montelionelab.chem.rpi.edu/dismeta/) and PONDR (https://www.pondr.com/). Protein domains were identified using HHpred (https://toolkit.tuebingen.mpg.de/tools/hhpred) (Zimmermann *et al*., 2018) , using default parameters, an E-value cutoff of 10^−5^, and searches against the Pfam and PDB databases. Protein structures were predicted using AlphaFold 3 (https://alphafoldserver.com/) (Abramson *et al*., 2024) and the resulting models were visualized and analyzed in UCSF ChimeraX v1.8 (Meng *et al*., 2023). DTH1 orthologs were identified using ProtBLAST (Zimmermann *et al*., 2018), using the DTH1 protein sequence lacking the coiled-coil domain (residues 141–911) as the query. Searches were performed with an E-value cutoff of 10^−5^ and the default BLOSUM62 scoring matrix. 11 hits were obtained. Only 6 were kept based on protein length and redundancy threshold fix at 90% and aligned in Jalview software using MAFFT v.7.

### Chlamydomonas strains and cultivation conditions

All Chlamydomonas strains WT (CC4533), *dth1-2* (LMJ.RY0402.135887) and *dth1-3* (LMJ.RY0402.198754) were maintained on agar plates containing TAP medium under continuous light at 25°C. Liquid cultures were grown in TAP liquid medium in conical glass flasks placed in incubators (Multitron, Infors HT) at 25°C with continuous illumination (80-100 µmol photons m^−2^ s^−1^) and agitation at 120 rpm. Cell concentration, size, and volume were measured using a Multisizer 4 instrument (Beckman Coulter). To initiate N deprivation, log-phase cultures were washed with nitrogen-free TAP medium (NH_4_Cl replaced by an equal molar amount of NaCl) (TAP-N) by centrifugation at 800 *g* for 5 min. After 48 h in TAP-N, cells were transferred to an N-containing minimum medium (MM) lacking acetate and incubated in the dark at 25°C. Under these conditions, cells were forced to mobilize stored carbon reserves as their sole source of carbon and energy.

### LD purification from Chlamydomonas

Five independent cultures of all three strains (CC4533, *dth1-2*, *dth1-3*) were cultivated and harvested at defined stages of LD dynamics: during LD formation (TAP-N, 48 h) and LD degradation (MM+N, 12 h). At each stage, LDs were isolated following the protocol described by Nguyen *et al*., 2011. Briefly, cell pellets were mechanically disrupted using a French press (Thermo Electron Aminco, USA) at 1500 psi, in Hepes-KOH buffer (25 mM, pH 7.5) supplemented with 0.6 M sucrose and protease inhibitor cocktail (Sigma-Aldrich, USA). The resulting lysate was gently layered into ultracentrifuge tubes containing the same buffer supplemented with 0.4 M sucrose to generate a sucrose density gradient, followed by centrifugation at 10,000 *g* for 30 min at 4°C using an Optima L-80 XP Ultracentrifuge rotor SW 32 Ti (Beckman Coulter). The floating lipid pad (∼3 mL) was carefully collected and subjected to the three sequential purification steps to remove proteins non-specifically associated with the LD fraction: (i) a detergent wash, with 0.1% Tween-20, (ii) an ionic wash with 2 M NaCl, and (iii) a chaotropic wash with 9 M urea. Purified LDs were then recovered through a final sucrose-gradient centrifugation step. A detailed step-by-step protocol is provided in **Figure S4**.

Two-thirds of the isolated LD fraction was used for proteomics analysis, while the remaining one-third was allocated for lipidomics analysis.

### Lipid extraction and analyses

#### Lipid extraction

Lipids from whole cells or isolated LDs were extracted using an isopropanol and methyl tert-butyl ether (MTBE)-based protocol adapted from (Légeret *et al*., 2016). Briefly, 10 million of cells or 500 µL of crude LD extracts were first boiled in 1 mL hot isopropanol supplemented with 0.01% (w/v) butylated hydroxytoluene (BHT). MTBE and aqueous NaCl 0,9% (w/v) were added in a ratio of isopropanol/MTBE/NaCl (1:3:1, v/v/v) to allow phase separation, vortexed vigorously, and centrifuged for 3 min at 3000 *g* at 4°C. The upper organic phase was collected in a separate clean glass tube, and MTBE extraction was repeated a second time by adding an additional 1 mL MTBE to the initial glass tube. The upper phase collection tube was evaporated under a gentle N_2_ stream, and the dried lipids were resuspended in 1 mL chloroform:methanol (2:1, v/v), supplemented with 10 µg TAG C15 and 10 µg PE 17:0 standards (prepared from 1 mg mL^−1^ stock solution in chloroform) to estimate the efficiency of polar/neutral lipid separation.

#### Polar/neutral lipid separation

Out of the 1 mL of extracted lipids, 200 µL were kept for total lipid analysis by GC-MS or LC-MS and 800 µL were used for polar/neutral lipid extraction. In brief, the 800 µL were evaporated, resuspended in 1 mL of acetonitrile:methanol:water (4.5:4.5:1, v/v/v) and vigorously shaken. The neutral fraction was collected by adding 2 mL of hexane to the mixture, vortexing, and centrifuging at 3,000 *g* for 3 min. The upper phase (i.e., the neutral fraction) was then collected in a clean glass tube. Hexane extraction was repeated twice (2 mL then 1 mL of hexane added). The lower phase was extracted with 3 mL MTBE, as it contained water and needed to be removed. After vortexing and centrifugation at 3,000 *g* for 3 min, the upper phase of MTBE extraction (i.e., the polar fraction) was collected in a new glass tube, evaporated under a gentle stream of N_2_ and resuspended in 100 µL of chloroform: methanol (2:1, v/v). 50 µL were kept for LC-MS analysis and 50 µL for GC-MS analysis.

#### Transmethylation of lipids

the total lipid extracts (100 µL), 50 µL of the polar lipid fraction and the whole neutral lipid fraction were converted to fatty acid methyl esters (FAMEs) by acid-catalyzed transmethylation. Briefly, lipid extracts were dried under a flow of N_2_ gas and resuspended in 1 mL of methanol with 5% sulfuric acid (v/v), supplemented with 10 µg of internal standard TAG19:0 (from a stock solution of 1 mg mL^-1^ in chloroform). The mixture was vortexed vigorously and heated at 85°C for 90 min. After a cooling phase, FAMEs were extracted with hexane, vortexed, and 1.5 mL of NaCl 0.9% (w/v) was added for phase separation. The samples were vortexed again vigorously, and centrifuged at 3,000 *g* for 3 min. The upper phase was collected in 0.5 mL vials for GC-MS analyses.

#### GC-MS analysis

GC-MS is coupled to a flame ionization detector and mass spectrometry (GC-FID-MS) on an Agilent 7890A GC and Agilent 5975C MS system from Agilent Technologies, Palo Alto, CA, USA. Separation of individual FAMEs was accomplished using a a Zebron 7HG-G007-11 (Phenomenex) polar capillary column (length 30 m, internal diameter 0.25 mm, and film thickness 0.25 µm) was used. The GC operating conditions were as follows: an injector and flame ionization detector temperature set at 240°C, and an oven temperature program starting at 60°C for 2 min, followed by a linear increase to 150°C at 20°C min^-1^. Subsequently, there was another increase at 6°C min^-1^ from 150°C to 240°C, and this temperature was maintained for 3 min. Samples were injected in spitless mode (1 min) at 250°C. The MS was run in full scan over 40 to 350 amu (electron impact ionization at 70 eV). The carrier gas (H2) flow rate was set at 1 mL min^-1^. Identification of FAMEs was based on retention time and mass spectrum, while quantification was carried out using the FID trace, as described by Siaut et al. (2011). The GC-MS results of FAMEs of total/polar/neutral lipids were utilized to normalize the quantity of polar lipid find in each sample by LC-MS.

#### LC-MS lipidomics analysis

For LC-MS analysis, 10 µg of PE34:0 standard was added as an internal standard to the remaining 50 µL of polar lipid in chloroform: methanol (2:1, v/v), and then the polar fraction was analyzed using a Thermo Vanquish UPLC system coupled with a Thermo Orbitrap Q Exactive mass spectrometer operated in data-dependent acquisition (DDA) mode. Briefly, the lipid mixtures (2 µL) were initially separated using a KinetexTM C18 2.1 × 150 mm 2.6 μm column (Phenomenex, Torrance, CA, USA) at a flow rate of 0.3 mL min-1, with the column temperature maintained at 45°C. The mobile phase consisted of solvent A (acetonitrile/water 60:40, v/v, containing 10 mM ammonium acetate) and solvent B (isopropanol/acetonitrile 90:10, v/v, containing 10 mM ammonium acetate). Lipids were separated using the following gradient of solvent B: the gradient started at 27% B for 0.5 min, increased linearly to 97% B over the next 24 min, was held constant during 2 min, returned linearly to 27% B and was held constant until the end of the run (32 min). The Q-Exactive mass spectrometer was used in negative ion mode with the following source parameters: sheath gas at 50 arbitrary units (AU); sweep gas at 2 AU; auxiliary gas at 15 AU; spray voltage at 2.5 kV; capillary temperature, 300°C; S-lens RF level, 50; scan range, m/z 200–1100. MS/MS spectra were acquired using automatic gain control target and stepped normalized collision energy of 25 eV, 30 eV and 40 eV. The acquired raw data were processed using Freestyle software (Thermo Fisher Scientific, Waltham, MA, USA). Lipid molecular species were identified based on accurate mass measurements and lipid fragmentation patterns. Relative quantification was accomplished using TraceFinder based on intensity values of extracted masses of previously identified lipids. Lipid molecular species were denoted as lipid class (total number of carbon atoms: total number of double bonds).

### Protein analyses

#### Protein extraction

Proteins were extracted from 900 µL of isolated LDs in HEPES 25 mM pH 7.5 by vortexing samples in 1% (v/v) SDS followed by incubation on ice for 30 min. Proteins were then precipitated by adding 9 vol of cold acetone (-20°C) to 1 vol of sample, and the mixture was incubated overnight at -20°C. Protein precipitates were collected by centrifugation at 18000 *g* for 10 min at 4°C. The supernatant was discarded, and residual acetone was evaporated. For electrophoresis, protein pellets were directly resuspended in 100 µL of Laemmli buffer containing 1% LDS and 100 mM DTT, heated at 70°C for 20 min and 15 µL were loaded onto 4-12% NuPAGE Mini gels (Invitrogen). The protein quantity was then adjusted and aligned based on whole-lane band quantitation on gel images with Fiji (v1.54f).

### Silver nitrate staining

The gel was first fixed for 30 min in a fixing solution containing 20 mL ethanol, 5 mL acetic acid, and 25 mL water, then 30 min in sensitization solution (15 mL ethanol, 3.4 g sodium acetate, 260 µL glutaraldehyde 25% (w/v), and 0.1 g sodium thiosulfate 5H_2_O in a 50 mL water solution). Gels were washed three times for 5 min in water and stained in a silver solution for 30 min (50 mg silver nitrate and 15 µL formaldehyde in 50 mL water). Protein bands were developed in a development solution (2.5 g sodium carbonate and 13 µL formaldehyde in 100 mL water) in two successive baths: the first bath was discarded once the solution turned grey, and the second bath was continued until the protein spots reached the desired intensity. Finally, the reaction was stopped by incubating the gel for 10 min in a solution containing 0.73 g EDTA-Na2 in 50 mL water, followed by a final rinse with water.

#### MS-based quantitative proteomic analysis of LDs

Proteins, solubilized in Laemmli buffer were stacked in the top of 4-12% NuPAGE gels (Invitrogen), stained with R-250 Coomassie Blue (BIORAD), before in-gel digestion using modified trypsin (sequencing purity, Promega), as previously described (Casabona *et al*., 2013)except that Tris(2-carboxyethyl)phosphine hydrochloride was used instead of dithiothreitol. The resulting peptides were analyzed by online nanoliquid chromatography coupled to MS/MS (Ultimate 3000 RSLCnano and Orbitrap Exploris 480, Thermo Fisher Scientific) operating in data-dependent acquisition mode. For this purpose, the peptides were sampled on a precolumn (300 μm x 5 mm PepMap C18, Thermo Scientific) and separated in a 75 μm x 250 mm C18 column (Aurora Generation 3, 1.7 μm, IonOpticks) using a 120 min acetonitrile gradient. The MS and MS/MS data were acquired by Xcalibur (Thermo Fisher Scientific).

Peptides and proteins were identified by Mascot (version 2.8.0, Matrix Science) through concomitant searches against *C. reinhardtii* phytozome database (version 5.6), the mitochondrion and chloroplast protein sequences downloaded from NCBI, and a homemade database containing the sequences of classical contaminant proteins found in proteomic analyses (e.g. human keratins, trypsin). Trypsin/P was chosen as the enzyme and two missed cleavages were allowed. Precursor and fragment mass error tolerances were set at respectively at 10 and 20 ppm. Peptide modifications allowed during the search were: Carbamidomethyl (C, fixed), Acetyl (Protein N-term, variable) and Oxidation (M, variable). The Proline software (Bouyssié *et al*., 2020, version 2.2.0) was used for the compilation, grouping and filtering of the results (conservation of rank 1 peptides, peptide length ≥ 6 amino acids, false discovery rate of peptide-spectrum-match identifications < 1% (Couté *et al*., 2020), and minimum of one specific peptide per identified protein group). Proline was then used to perform a MS1-based label-free quantification of the identified protein groups based on razor and specific peptides.

Statistical analyses were conducted using Prostar (version 1.30.5, Wieczorek *et al*., 2017). Proteins identified in the contaminant database and proteins identified by MS/MS in less than three replicates of one condition and quantified in less than five replicates of one condition were discarded. After log2 transformation, abundance values were normalized using the variance stabilizing normalization (vsn) method, before missing value imputation (SLSA algorithm for partially observed values in the condition and DetQuantile algorithm for totally absent values in the condition). Statistical testing was conducted with limma, whereby differentially expressed proteins were selected using a log2(Fold Change) cut-off of 1 and a *p*-value cut-off of 0.01, allowing to reach false discovery rates inferior to 5.3 % according to the Benjamini-Hochberg estimator. Proteins found differentially abundant but identified by MS/MS in less than three replicates and detected in less than five replicates, in the condition in which they were found to be more abundant were manually invalidated (*p*-value = 1).

For each protein in each sample, niBAQ values were calculated by normalizing the extracted iBAQ (Schwanhäusser *et al*., 2011) values by the sum of all iBAQ values in the sample. A global niBAQ value for each protein in each condition was obtained by summing the five niBAQ values and dividing by five.

### Microscopy

#### Confocal imaging

All images were acquired using a Carl Zeiss LSM980 microscope equipped with a 63× oil immersion objective. BODIPY 505/515 fluorescence was excited at 488 nm and emission was detected between 495 – 530 nm, while AC202 (Harchouni *et al*., 2018) fluorescence was excited at 405 nm and emission was detected between 450 – 465 nm. The emission of the autofluorescence of chlorophyll was detected between 650 – 680 nm. All images represent a z-stack with a 0.2 µm step size, covering the cell from top to bottom. All fluorescence signals were analyzed using Fiji (v1.54f).

#### Quantification of LD size and number

LD size and number were calculated using Fiji. Briefly, the BODIPY or AC202 channel was used to create a binary mask for LDs, and the transmitted light channel was used to create a binary mask for cells. For each LD, area and centroid were calculated, and each LD was attributed to a cell based on the centroid position relative to the binary mask of cells. For each cell, the total number of LDs was calculated. Data output was then visualized using GraphPad Prism (v11).

#### Transmission electron microscope (TEM)

CC4533, *dth1-2* and *dth1-3* were fixed in suspension with 2.5 % glutaraldehyde and 0.1 M PBS (pH 7.2) at different time points: after 48 h of N starvation (NR0) and 6 h after N-resupply (NR6). Aliquots were post-fixed with reduced osmium tetroxide/potassium ferrocyanide (1 % OsO4 / 1.5 % K3Fe(CN)6), then treated with 1 % tannic acid in 0.1 M PBS, infiltrated with an ethanol/Quetol 651 resin (Electron Microscopy Sciences, USA) resin mixture, and embedded in Quetol. 90-nm-thick sections were cut with a diamond knife in a Leica UC6 ultramicrotome. The sections were collected on carbon-coated copper grids, post-stained with uranyl acetate and lead citrate, and observed with a JEOL JEM-2100-Plus microscope operating at an accelerating voltage of 200 kV. Images were recorded with a Gatan Rio 16 camera.

## Statistical analysis

Statistical analyses were performed using GraphPad Prism (GraphPad Software) and R (version 2024.12.0). Data normality was assessed using the Shapiro–Wilk test. As the data did not follow a normal distribution, non-parametric statistical tests were used. The Wilcoxon rank-sum test was used for comparisons between two independent groups, while the Kruskal–Wallis test followed by Dunn’s post-hoc test was used for comparisons involving more than two groups. P-values were adjusted for multiple comparisons using the Benjamini–Hochberg false discovery rate (FDR) correction. For all analyses, statistical significance (α=0.05) according to *p* values is indicated by asterisk (* *p* ≤ 0.05; ** *p* ≤ 0.01; *** *p* ≤ 0.001 and **** *p* ≤ 0.0001).

## Author contributions

Y.L.B and C.B. conceived this study. C.B. performed most of the experiments under the supervision of B.L. for lipid analysis, S.C. for LD purification, M.S. for protein sequence analysis, and M.B. for confocal microscopy. J.L.P. performed TEM analysis. A.T. and Y.C. performed MS-based proteomic analyses. C.B. and Y.L.B. wrote the manuscript with inputs from all co-authors. All authors approved the manuscript.

## Acknowledgements

The use of the HelioBiotec and Zoom platforms of the BIAM institute is acknowledged. We also thank the NanoBio-ICMG Platform (UAR 2607, Grenoble) for granting access to the electron microscopy facility, and Christine Lancelon-Pin (CERMAV) for the preparation of the ultrathin sections for the TEM analysis.

## Data and material availability

All biological material described in this study is available upon request. The mass spectrometry proteomics data have been deposited to the ProteomeXchange Consortium via the PRIDE (Perez-Riverol *et al*., 2022) partner repository with the dataset identifier PXD085091 and 10.6019/PXD085091.

## Supplemental figures

**Figure S1. Multiple sequence alignment of DTH1 with orthologs from green algae.**

The sequences of CrDTH1 and its orthologs were aligned using MAFFT program in Jalview software and only the C-terminal part of the alignment is shown here. Secondary structure elements predicted by AlphaFold3 are shown above the alignment (red, alpha-helices; black, beta-sheet). Residues that are identical across the sequences appear with a red background, while positions showing conservative amino acid substitutions are marked in blue. Abbreviations: Astgub, *Astrephomene gubernaculifera*; Chleu, *Chlamydomonas eustigma*, CrDTH1, Delayed in Triacylglycerol Hydrolysis 1 (*Chlamydomonas reinhardtii*); Edade, *Edaphochlamys debaryana*; Volafr, *Volvox africanus*; Volca, *Volvox carteri_f_nagariensis*.

**Figure S2. Polar lipid composition of wild-type (CC-4533) and *dth1* mutants under N-starvation and N-resupply.**

**A.** Experimental workflow. 10 million cells were harvested after 48 h of N starvation (NR0), or after 48 h of N starvation followed by 5 h of N resupply (NR5), and then polar and neutral lipids were separated from total extracts.

**B.** Polar lipid class composition in whole cells CC-4533 and *dth1* mutants under NR0 and NR5 conditions. Lipid species peak areas were summed per class for each replicate.

**C.** Polar lipid species composition in wild-type and *dth1* mutants under NR0 and NR5. Species < 0.5% of total response were omitted. For panel (B) and (C), data are mean ± SD from 6–10 replicates; significance was assessed by Kruskal–Wallis and Dunn’s test with FDR correction (*\*p* < 0.05, *\*\*p* < 0.01).

Abbreviations: DGDG, digalactosyldiacylglycerol; DGTS, diacylglyceryl-*N,N,N*-trimethylhomoserine; MGDG, monogalactosyldiacylglycerol; PE, phosphatidylethanolamine; PG, phosphatidylglycerol; PI, phosphatidylinositol; SQDG, sulfoquinovosyldiacylglycerol.

**Figure S3. TEM observations of WT and *dth1* mutants.**

**A-C.** TEM images of WT in NR6 (A and C) and *dth1-2* in NR6 (B) where we observed LDs moving out into the periplasmic space of the cell for some unknown reasons (blue arrow) and they are degraded in that subcellular compartment (red arrow) by some unknown excreted TAG lipases. We recurrently observed mitochondria, making contact with LDs on both sides of LDs that are moving outward.

**D-E**. mLDs (orange arrows) observed in the periplasm of CC-4533 (C), and *dth1-3* (D) in NR6 conditions.

**F.** Autophagosome found in the periplasm containing an LD with mLDs around it in *dth1-3* under NR6.

**G.** Vesicle exocytosis containing LD being degraded and mLDs. We can not say whether this is a lytic vacuole or an autophagosome.

**H.** mLDs observed around starch granules.

**I.** mLDs observed in flagella.

**Figure S4. Lipid droplet extraction and purification protocol and proteins extraction.**

**A.** Complete procedure for LD extraction and purification, based on sucrose-gradient floatation coupled with several steps of washes to remove contaminants from LD monolayer.

**B.** Silver-stained SDS-PAGE of LD proteins extracted during TAG accumulation (NR0) and TAG degradation (NR12).

Equal amounts of total protein were loaded based on adjusted whole-lane band quantitation on gel images. Five replicates from each genotype were analyzed. MLDP: major lipid droplet protein; NR: nitrogen resupply.

**Supplemental tables**

**Supplemental table S1: MS-based quantitative proteomic analysis of lipid droplets purified from CC4533 and *dth1* mutant cells exposed to nitrogen starvation (NR0) and resupply (NR12).**

