## Supplementary figures and images for "DTH1-dependent remodeling of the lipid and protein environment of lipid droplets during nutrient recovery in Chlamydomonas"

### Supplemental figure S1

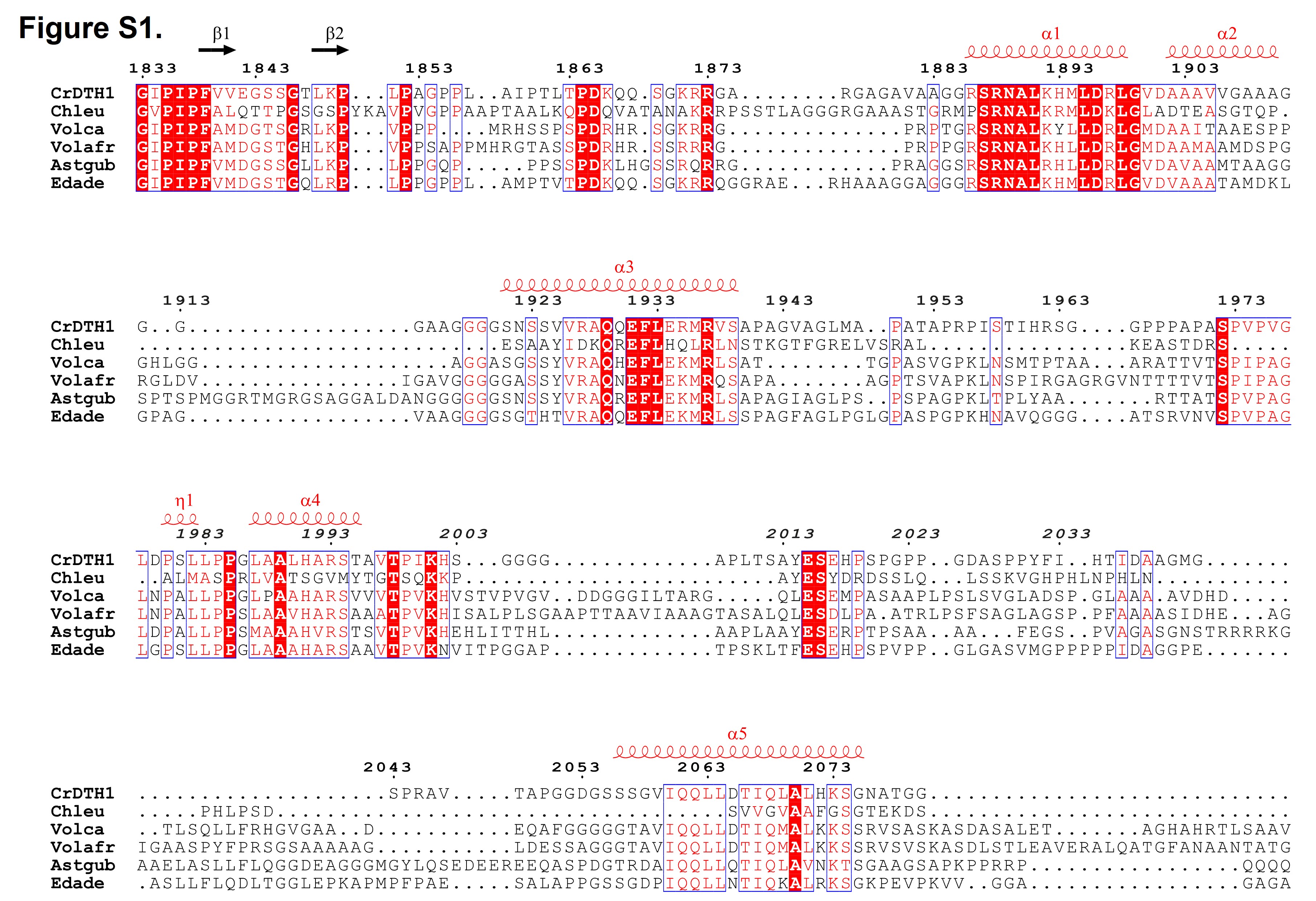

### Supplemental figure S2

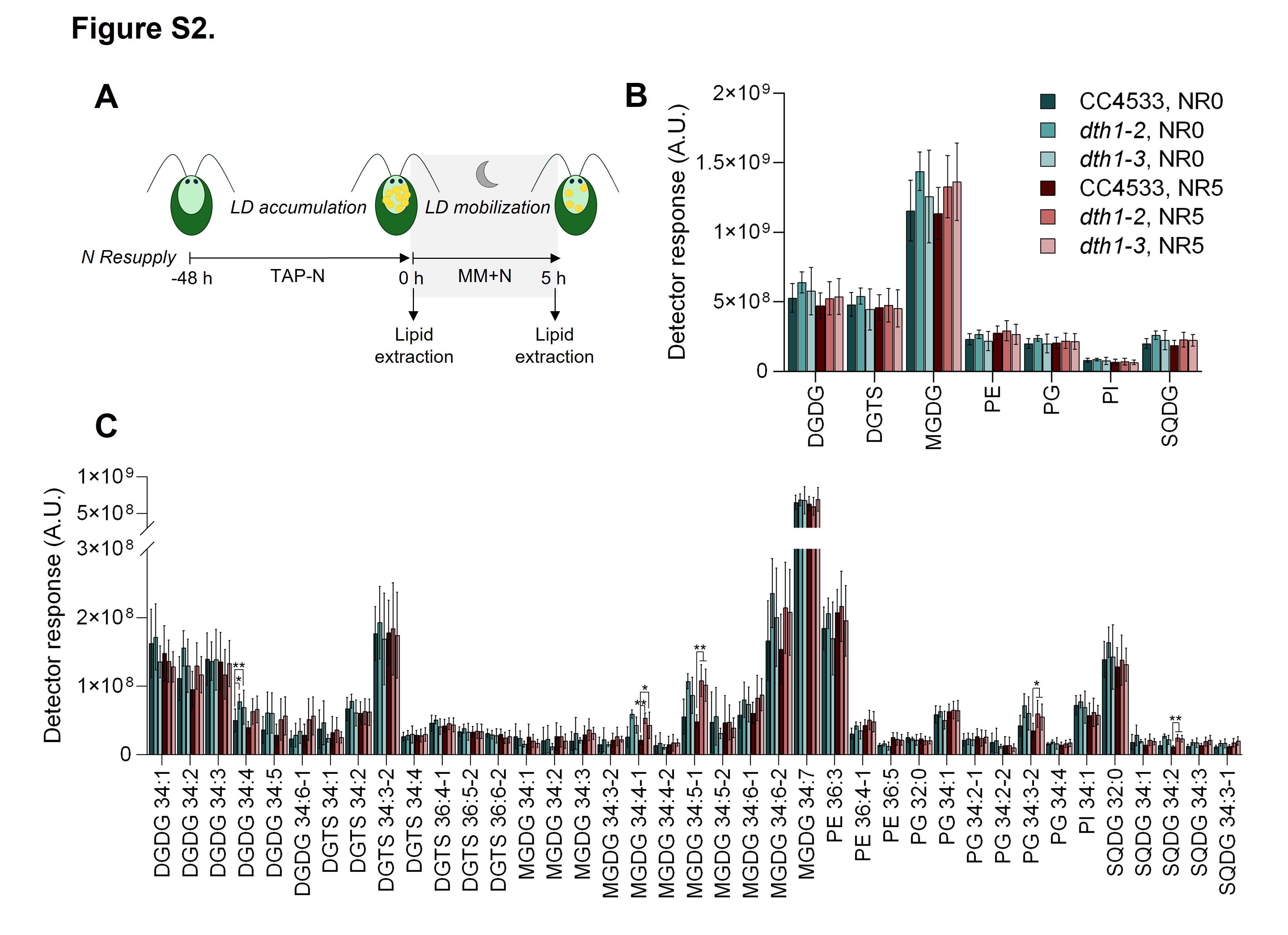

### Supplemental figure S3

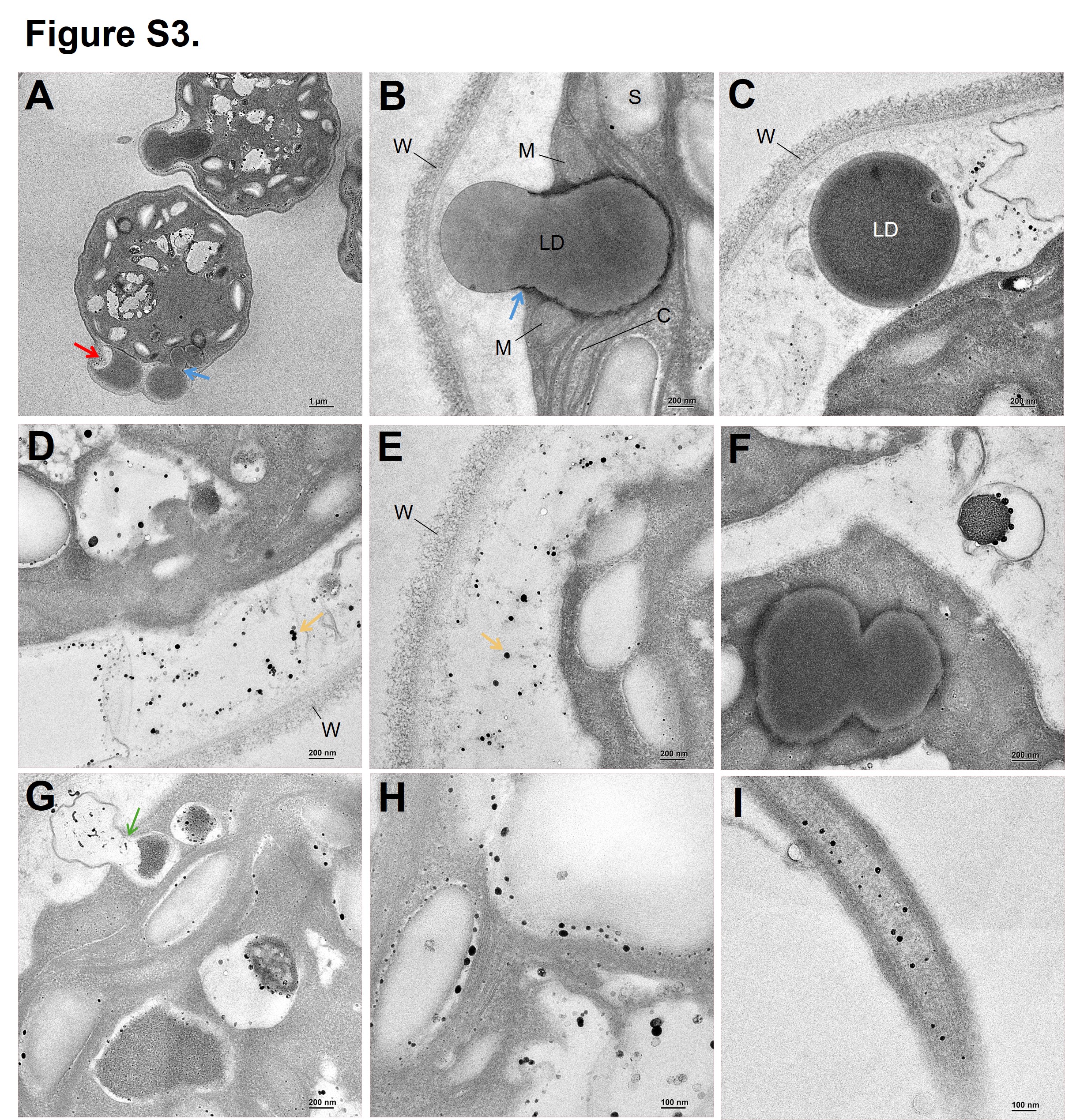

### Supplemental figure S4

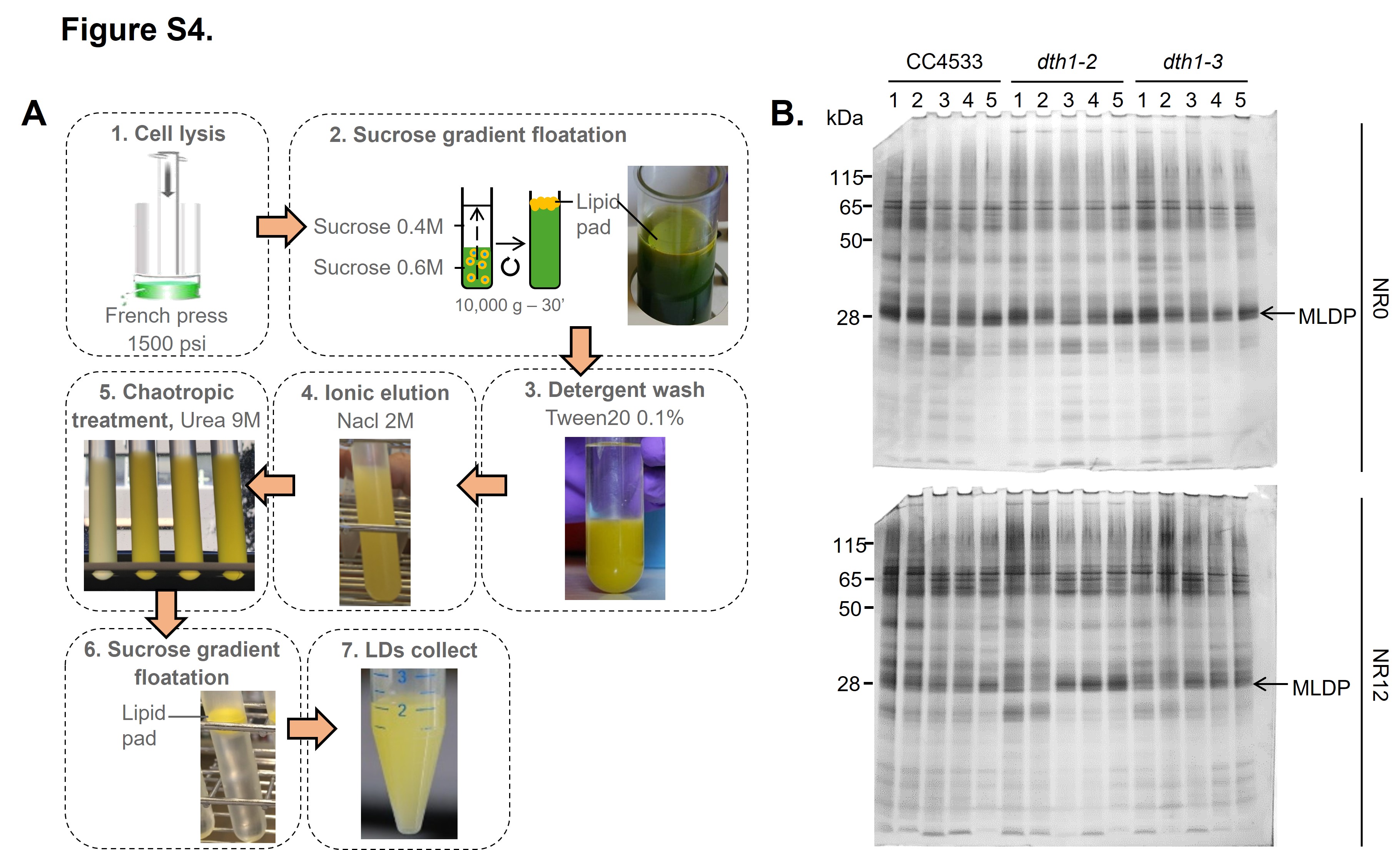
